# Enhancing anti-EGFRvIII CAR T cell therapy against glioblastoma with a paracrine SIRPγ-derived CD47 blocker

**DOI:** 10.1101/2023.08.31.555122

**Authors:** Tomás A. Martins, Nazanin Tatari, Deniz Kaymak, Sabrina Hogan, Ewelina M. Bartoszek, Ronja Wieboldt, Marie-Françoise Ritz, Alicia Buck, Marta McDaid, Alexandra Gerber, Aisha Beshirova, Manina M. Etter, Anja Heider, Tala Shekarian, Hayget Mohamed, Philip Schmassmann, Ines Abel, Luigi Mariani, Raphael Guzman, Jean-Louis Boulay, Berend Snijder, Tobias Weiss, Heinz Läubli, Gregor Hutter

**Affiliations:** Brain Tumor Immunotherapy and Biology Laboratory, Department of Biomedicine, University of Basel, Basel, Switzerland; Microscopy Core Facility, Department of Biomedicine, University of Basel, Basel, Switzerland; Cancer Immunotherapy Laboratory, Department of Biomedicine, University of Basel, Basel, Switzerland; Department of Neurology, Clinical Neuroscience Center, University Hospital Zurich and University of Zurich, Zurich, Switzerland; Institute of Molecular Systems Biology, ETH Zurich, Zurich, Switzerland; Experimental Immunology Laboratory, Department of Biomedicine, University of Basel, Basel, Switzerland; Department of Neurosurgery, University Hospital Basel, Basel, Switzerland; Immunology Laboratory, Swiss Institute of Allergy and Asthma Research, University of Zurich, Davos Wolfgang, Switzerland; Division of Oncology, University Hospital Basel, Basel, Switzerland; Department of Surgery, University Hospital Basel, Switzerland

**Keywords:** GBM, EGFRvIII, CAR T cell therapy, CD47, GAM, microglia modulation, SGRP, immunotherapy

## Abstract

A major challenge for chimeric antigen receptor (CAR) T cell therapy against glioblastoma (GBM) is its immunosuppressive tumor microenvironment (TME), which is densely populated and supported by protumoral glioma-associated microglia and macrophages (GAMs). Targeting of CD47, a “don’t-eat-me” signal overexpressed by tumor cells, disrupts the CD47-SIRPα axis and induces GAM phagocytic function. However, antibody-mediated CD47 blockade monotherapy is associated with toxicity and low bioavailability in solid tumors. To overcome these limitations, we combined local CAR T cell therapy with paracrine GAM modulation for more effective elimination of GBM. To this end, we engineered a new CAR T cell against epidermal growth factor receptor variant III (EGFRvIII) that constitutively secretes a SIRPγ-related protein (SGRP) with high affinity to CD47. Anti-EGFRvIII-SGRP CAR T cells eliminated EGFRvIII^+^ GBM in a dose-dependent manner *in vitro* and eradicated orthotopically xenografted EGFRvIII-mosaic GBM by locoregional application *in vivo.* This resulted in significant tumor-free long-term survival, followed by partial tumor control upon tumor re-challenge. The combination of anti-CD47 antibodies with anti-EGFRvIII CAR T cells failed to achieve a similar therapeutic effect, underscoring the importance of sustained paracrine GAM modulation. Multidimensional brain immunofluorescence microscopy and in-depth spectral flow cytometry on GBM-xenografted brains showed that anti-EGFRvIII-SGRP CAR T cells accelerated GBM clearance, increased CD68^+^ cell trafficking to tumor scar sites, and induced myeloid-mediated tumor cell uptake. Additionally, in a peripheral lymphoma mouse xenograft model, anti-CD19-SGRP CAR T cells had superior efficacy compared to conventional anti-CD19 CAR T cells. Validation on human GBM explants revealed that anti-EGFRvIII-SGRP CAR T cells had similar tumor-killing capacity to anti-EGFRvIII CAR monotherapy, but showed a slight improvement in maintenance of tumor-infiltrated CD14^+^ myeloid cells. Thus, local anti-EGFRvIII-SGRP CAR T cell therapy combines the potent antitumor effect of engineered T cells with the modulation of the surrounding innate immune TME, resulting in the additive elimination of bystander EGFRvIII^-^ tumor cells in a manner that overcomes major mechanisms of CAR T cell therapy resistance, including tumor innate immune suppression and antigen escape.

## Introduction

Glioblastoma (GBM) is the most aggressive malignant primary brain tumor in adults. The infiltrative nature of GBM, and its localization near eloquent areas of the brain, often make complete surgical resection unachievable. Moreover, chemo- and radiotherapy regimens are not curative, invariably leading to recurrent disease. The survival time for patients with GBM is around 15 months, underscoring an urgent need for a significant breakthrough in effective medical treatment^1–3^.

Chimeric antigen receptor (CAR) T cell-based immunotherapies have had remarkable outcomes in the clinical treatment of hematological malignancies, yet the development of effective CAR T cell therapies against solid tumors remains challenging^4,5^. A critical limitation of CAR T cell therapy against GBM is the scarcity of known tumor-specific surface antigens and their heterogeneous expression pattern^6^. One of the most well-studied target antigens in GBM is epidermal growth factor receptor variant III (EGFRvIII), a tumor-specific mutated form of epidermal growth factor receptor (EGFR) expressed in approximately 40% of GBM cases^7,8^. Although strictly expressed on tumor cells, EGFRvIII mutations arise with concomitant EGFR amplification during clonal evolution events in GBM development, resulting in EGFRvIII-mosaic tumors^8,9^. A clinical trial of anti-EGFRvIII CAR T cells against recurrent GBM (NCT01454596) showed safety and transient efficacy but failed to produce long-lasting treatment responses due to adaptive resistance and antigen escape^10^. Conversely, targeting more homogeneously expressed GBM-associated antigens is controversial due to the potential risk of on-target/off-tumor cross-reactions leading to toxicity^11,12^.

Immune checkpoint inhibitors have recently shown promising responses against solid tumors^13^. However, the highly immunosuppressive tumor microenvironment (TME) of GBM severely limits the efficacy of immune checkpoint blockade (ICB)^14^. Thus, understanding the complex context-dependent interactions of GBM with the surrounding TME is crucial for the effective targeting of these tumors using immunotherapeutic approaches^15^.

The most important and dominant cell populations in the GBM immune compartment are pro-inflammatory and proliferative brain-resident microglia, peripheral monocyte-derived macrophages, polymorphonuclear myeloid-derived suppressor cells and T_regs_^16,17^. Notably, microglia and macrophages are professional phagocytic cells that play a key role in brain innate immune surveillance. In GBM, they are converted into glioma-associated macrophages and microglia (GAMs) that support tumor growth through the release of various cytokines, chemokines, and growth factors^11,18,19^. The phagocytic activity of GAMs is regulated through, among others, the CD47-SIRPα phagocytosis axis, whereby SIRPα, expressed on the surface of phagocytes, interacts with the ubiquitously expressed CD47 transmembrane protein, thereby inhibiting phagocytosis^20,21^. Hence, CD47 is an innate immune checkpoint co-opted by tumor cells leading to immune evasion through their reduced recognition by phagocytes^11,22^. It has been shown that CD47 blockade rescues GAM phagocytic function in GBM-bearing mice resulting in a strong antitumoral response *in vivo*^23–25^. However, clinical studies of systemic anti-CD47 monotherapy have only recently begun to assess efficacy against solid tumors, showing promising clinical activity^26,27^. Published results from these trials suggest overall safety and considerable activity but also report low bioavailability within the tumor and a high level of treatment-associated toxicity^28^.

Myeloid immunosuppression and clonal heterogeneity are the main hurdles for effective anti-cancer immunotherapies against GBM^29^. To synergistically target heterogeneous GBM and the immunosuppressive TME, we propose a combinatorial approach of intratumoral (i.t.) CAR T cell therapy and GAM modulation for the targeting of antigen-expressing and bystander tumor cells, yielding complete GBM clearance. Here we show that the combination of tumor-targeting CAR T cells with the constitutive secretion of a paracrine SIRPγ-related protein (SGRP) dramatically improved the elimination of EGFRvIII-mosaic GBM *in vivo* in orthotopic GBM and peripheral lymphoma xenograft mouse models. The proof-of-concept of the therapeutic strategy shown here could lead the way for improved CAR T cell therapies in GBM and other solid tumor types.

## Materials and Methods

### SGRP engineering for production and release by CAR T cells

To generate SGRP, 9 amino acid (AA) substitutions were made to the endogenous human SIRPγ binding domain (hSIRPγ-V1) AA sequence, as previously described by Ring and colleagues^30^ (**Fig. S1A**). A human IL-2 signal peptide (IL2sig) sequence was added to the N-terminus of SGRP to allow its secretion by T cells to the extracellular space (**Fig. 1C**). The SGRP AA sequence was codon-optimized using GenSmart Codon Optimization (GenScript Biotech, USA) with ‘Human T-cell’ set as the expression host organism. The resulting optimized AA sequence was then reverse-translated into a nucleotide (nt) sequence. All post-optimization AA and nt sequences are listed in **Table S1**.

**Figure 1.**
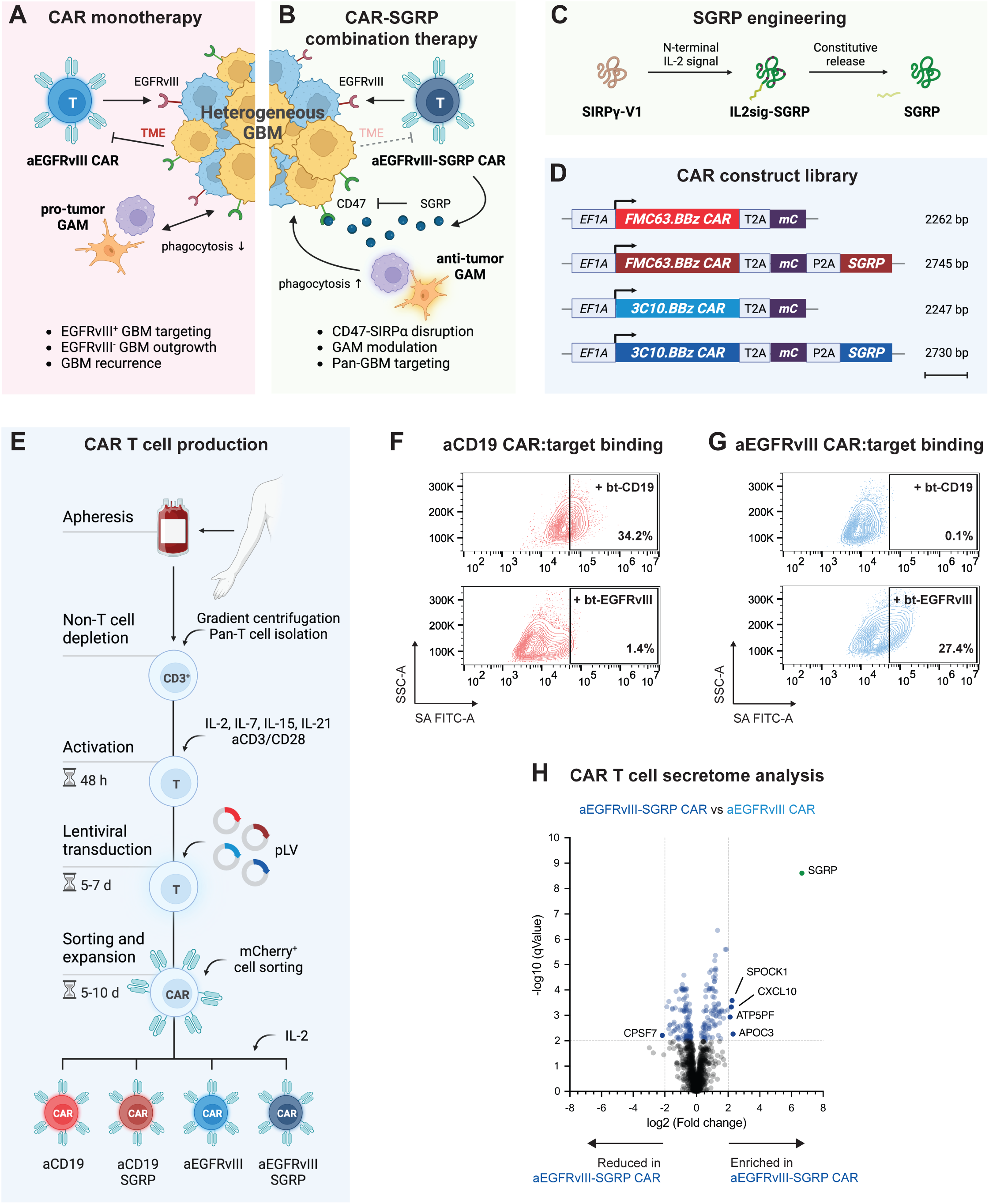
Anti-EGFRvIII-SGRP CAR T cells constitutively secrete SGRP, enabling SIRPα-CD47 axis disruption. **A,** Proposed mechanism of action of conventional aEGFRvIII CAR T cell monotherapy. **B,** aEGFRvIII-SGRP CAR T cell combination therapy whereby SGRP-mediated CD47 blockade induces phagocytic modulation of GAMs in the context of EGFRvIII-heterogenous GBM and its immunosuppressive TME. **C,** Outline of the SGRP engineering strategy including specific AA substitutions to the endogenous human SIRPγ-V1 sequence and addition of an N-terminal IL-2 signal sequence (IL2sig) leading to constitutive SGRP secretion. Full CAR sequences are listed in **Table S2**. **D,** Polycistronic lentiviral constructs encoding mCherry (mC)-labeled aCD19 CAR or aEGFRvIII CAR under the control of *EF1A* promoter +/- SGRP secretion. **E,** Workflow of CAR T cell production from HD PBMC-derived CD3^+^ T cells applied throughout the study. **F,** and **G,** Representative plots of CAR:target protein binding by aCD19 CAR T cells (**F**) or aEGFRvIII CAR T cells (**G**) assessed by FC using streptavidin (SA)-FITC labeling of CAR-bound biotinylated (bt)-CD19 (top plots) or bt-EGFRvIII (bottom plots); n = 2 HDs assessed per CAR. **H,** Differentially secreted proteins in CAR T cell-conditioned media, detected by LC–MS, of 24 h-rested, antigen-naïve aEGFRvIII-SGRP CAR vs aEGFRvIII CAR, highlighting the presence of SGRP (green dot) exclusively in aEGFRvIII-SGRP CAR T cell-conditioned media; n = 2 HDs. Mean SGRP expression: -log10^qValue^/log2^fold-change^ = 8.60/6.65.

### CD47-SIRP/SGRP protein interaction modeling

Protein structures of SGRP, human SIRPα binding domain (hSIRPα-V1), murine SIRPα binding domain (mSIRPα-V1) and human CD47 (hCD47) were predicted by AlphaFold^31,32^, using the source code, trained weights and inference script available under an open-source license: https://github.com/deepmind/alphafold. AA sequences of SGRP, hSIRPα-V1, mSIRPα-V1 and hCD47 were entered and folded using the multimer model. Sequences were superimposed against a genetic database to generate multiple sequence alignment statistics. Predictions ran on ‘relax mode without GPU’ and generated 3D interaction models for SGRP and hCD47, hSIRPα-V1 and hCD47, or mSIRPα-V1 and hCD47 (**Fig. S1B**). Predicted local distance difference tests (pLDDT) were calculated to evaluate local distance differences of all atoms in each model and validate stereochemical plausibility (data not shown). All AA and nt sequences are listed in **Table S1**.

### CAR construct and lentiviral expression vector design

For human T cell transduction, replication-defective lentiviruses were produced using a second-generation lentiviral system with transfer plasmids encoding a 3C10.BBz^33,34^ (anti-EGFRvIII) or FMC63.BBz^35,36^ (anti-CD19) CAR, an mCherry fluorescence reporter protein and, in some iterations, SGRP^30,37^. The CAR structure consisted of a CD8α leader, a single-chain variable fragment (scFv), CD8α hinge and transmembrane domains, a 4-1BB costimulatory domain and a CD3ζ signaling domain. Transgene expression was driven by the *EF1A* promoter and polyprotein sequences were cleaved by T2A or P2A peptides. All sequences were assembled with Vector Design Studio (VectorBuilder, USA). The vectors included an ampicillin resistance gene for positive selection of transformed *E. coli*. The lentiviral plasmids were purchased as bacterial glycerol stocks from VectorBuilder (VectorBuilder, USA). CAR constructs and lentiviral vector maps are illustrated in **Fig. 1D** and **Fig. S1C**, respectively. CAR construct and CAR domain sequences are listed in **Table S2**.

### Lentivirus production

Plasmid DNA was extracted with a QIAprep Spin Miniprep Kit (#27104, QIAGEN, Netherlands) from bacterial cultures grown overnight in the presence of 100 µg/mL ampicillin (#A5354, Sigma-Aldrich, USA). Lentiviral particles were generated by co-delivery of a transfer plasmid, a plasmid encoding a VSV-G envelope (pMD2.G; #12259, Addgene, USA) and an empty backbone plasmid (psPAX2; #12260, Addgene, USA) into HEK293T cells. Cells were maintained in DMEM (#11995065, Gibco, USA) supplemented with 10% inactivated fetal bovine serum (FBS; #P30-3302, PAN-Biotech, Germany), 1% pen strep (#15140-122, Gibco, USA) and 2 mM GlutaMAX-I (#35050-038, Gibco, USA). They were cultured as adherent monolayers, at 37°C, in a 5% CO_2_ atmosphere and regularly sub-cultured when reaching approximately 70-80% confluence. For the transfections, 5 × 10^5^ HEK293T cells were seeded per well in a 6-well plate and rested for 24 h. Growth media were replaced with antibiotic-free growth media, DNA plasmids were complexed with polyethyleneimine (PEI; #408727, Sigma-Aldrich, USA) for 15 min and added dropwise to the cells, followed by a 48 h incubation. The viral supernatants were collected and cleared from cells and debris by centrifugation at 500 × g for 10 min, followed by filtration through a 0.45 µm polyethersulfone filter (#SLHPM33RS, MilliporeSigma, USA). Virus particles were precipitated with Lenti-X Concentrator (#631232, Takara Bio, Japan), suspended in phosphate buffer saline (PBS; #D8537, Sigma-Aldrich, USA), quantified with Lenti-X GoStix Plus (#631280, Takara Bio, Japan), aliquoted and stored at -80°C.

### Healthy-donor T cell isolation

Peripheral blood leukocytes from healthy donors (HDs) were obtained from the Blood Donation Center of the University Hospital Basel, Switzerland, after informed consent was obtained from all participants before blood collection. Peripheral blood mononuclear cells (PBMCs) were isolated by Ficoll Paque-PLUS (#GE17-1440-02, Cytiva, Germany) and density centrifugation. After up to 2 rounds of ACK-lysis (#A10492-01, Gibco, USA) for removal of erythrocytes, PBMCs were washed with PBS. CD3^+^ T cells were magnetically separated by negative selection with a Human Pan T cell isolation Kit (#130-096-535, Miltenyi Biotec, Germany) and stored long-term in Bambanker serum-free cell freezing medium (#BB01, GC Lymphotec, Japan) in liquid nitrogen (LN_2_).

### CAR T cell production

CAR T cells were produced from HD PBMCs, depleted for non-T cells and frozen in batches. CAR T cells were freshly produced for every experiment from batches of frozen HD T cells. Upon thawing, T cells were washed with PBS and rested in X-VIVO 15 (#BE02-060F, Lonza, Switzerland) at a density of 1 × 10^6^ cells per mL at 37°C, in a 5% CO_2_ atmosphere. After 24 h, the T cells were activated in X-VIVO 15 containing 150 U/mL of human IL-2 (#Ro 23-6019, Roche, Switzerland), 10 ng/mL of recombinant IL-7 (#200-07, PeproTech, USA), 10 ng/mL of recombinant IL-15 (#200-15, PeproTech, USA), 20 ng/mL of recombinant IL-21 (#200-21, PeproTech, USA) and Dynabeads human T-activator CD3/CD28 (aCD3/CD28; #11131D, Gibco, USA) in a 1:1 cell-bead ratio^38^. After 48 h, aCD3/CD28 beads were magnetically removed and T cells were resuspended in X-VIVO 15 with 5 µg/mL polybrene (#TR-1003-G, Sigma-Aldrich, USA) at a density of 3 × 10^6^ cells per mL. Lentiviral suspensions were added to the T cells at different multiplicity of infection (MOI) ratios and spinfected at 2500 rpm, at 30°C for 90 min. After spinfection, the T cells were washed and maintained at a density of 1 × 10^6^ cells per mL in X-VIVO 15 containing 500 U/mL IL-2 for 5-7 days. After this post-transduction cell expansion, T cells were sorted for mCherry expression using a BD FACSMelody Cell Sorter (BD Biosciences, USA). After sorting, the CAR T cell cultures were expanded in X-VIVO 15 containing 500 U/mL IL-2 and kept at a density of 1 × 10^6^ cells per mL by adjusting the cell density every 2-3 days based on automated cell counting with a LUNA-FL Dual Fluorescence Cell Counter (Logos Biosystems, South Korea) for 5-10 days until used in downstream assays. A schematic of CAR T cell production from HD T cells is illustrated in **Fig. 1E**.

### CAR-biotinylated target protein binding assay

CAR T cell viability and count were assessed by Trypan blue exclusion. Cells were washed with PBS and seeded into a 96-well plate at a density of 2 × 10^5^ live cells per well. Cells were immediately stained with a Zombie NIR Viability kit (#423106, BioLegend, USA) diluted 1:5000 in PBS for 20 min in the dark at RT. After viability staining, the cells were washed with autoMACS Running Buffer (#130-091-221, Miltenyi Biotec, Germany) and then resuspended in 100 µL per well of 10 µg/mL dilutions of biotinylated CD19, EGFR or EGFRvIII proteins (#CD9-H82E9, #EGR-H82E3 and #EGR-H82E0, ACROBiosystems, USA) for 1 h in the dark at 4°C. After CAR-target exposure, the cells were washed with autoMACS Running Buffer and stained with 100 µL per well of FITC Streptavidin (SA; #405202, BioLegend, USA) diluted 1:50 in autoMACS Running Buffer, for 1 h in the dark at 4°C. Afterward, the cells were washed 3 times with autoMACS Running Buffer and resuspended in 100 µL of autoMACS Running Buffer. Samples were acquired with a CytoFLEX Flow Cytometer (Beckman Coulter, USA) and data were analyzed using FlowJo v10 Software (BD Biosciences, USA).

### CAR T cell secretome analysis

Expanded aEGFRvIII CAR and aEGFRvIII-SGRP CAR cultures were rested in X VIVO medium without additional supplements for 24 h. The following day, cell viability and count were assessed by Trypan blue exclusion, after which cells were washed in PBS, resuspended in RPMI at a density of 1 × 10^6^ live cells per mL and incubated for another 24 h at 37°C. Afterward, cultures were centrifuged at 300 × g for 5 min, supernatants were collected, passed through 0.22 µm filters and stored at -20°C. Proteins present in cell culture supernatants were precipitated by adding 1 volume of 100% (w/v) trichloroacetic acid (TCA) to 4 volumes of sample in microcentrifuge tubes. After 10 min incubation at 4°C, the samples were centrifuged at 14000 rpm for 5 min and supernatants were discarded. Undisturbed protein pellets were gently washed with 200 µL of pre-cooled acetone and then washed a second time with 200 µL of cold acetone followed by 10 sec vortexing. Samples were centrifuged at 14000 rpm for 5 min, supernatants were discarded and 20 µL of GuA digestion buffer (6 M guanidium hydrochloride, 100 mM ammonium bicarbonate (AB; pH = 8.3), 10 mM tris(2-carboxyethyl)phosphine (TCEP), 15 mM chloroacetamide was added. After 10 min incubation at 95°C with gentle agitation, samples were sonicated in a Bioruptor Pico (Diagenode, Belgium) for 10 cycles of 30 sec on and 30 sec off. Diluted GuA digestion buffer in the samples to 0.5 M by adding 100 mM AB. Trypsin (#V5280, Promega, USA) was then added to the samples in a 1:50 trypsin:sample ratio and incubated overnight at 37°C. The following day, samples were acidified to pH < 2 with 50 µL of 20% TCA. C18-columns were conditioned with 200 µL of acetonitrile and equilibrated 2 times with 200 µL C18-buffer A. Samples were then loaded into the C18-columns and centrifuged at 110 × g for 1 min. Flow-throughs were reloaded onto the columns and centrifuged again for 2 min. Columns were washed 3 times with 200 µL of C18-buffer C and bond peptides were eluted with 2 times 100 µL C18-buffer B into 2 mL microcentrifuge tubes. Finally, eluted peptide mixtures were concentrated to dryness under a vacuum. Samples were rehydrated and loaded into a spectrometer. Peptide spectra were analyzed using differential expression analysis, comparing aEGFRvIII-SGRP CAR and aEGFRvIII CAR samples from 2 HDs. A full list of detected proteins and their relative abundance is provided in **Table S3**.

### Cell lines and cell culture

BS153, U87, U251 and U251vIII are human glioma cell lines. BS153 cells were maintained in DMEM (#10938025, Gibco, USA) supplemented with 10% inactivated FBS, 1% pen strep, 2 mM GlutaMAX-I and 1 mM sodium pyruvate (#S8636, Sigma-Aldrich, USA). U87, U251 and U251vIII cells were maintained in MEM (#M4655, Sigma-Aldrich, USA) supplemented with 10% inactivated FBS, 1% pen strep, 1X MEM NEAA (#11140-035, Gibco, USA), 2 mM GlutaMAX-I and 1 mM sodium pyruvate. Raji is a lymphoma cell line cultured in DMEM supplemented with 10% inactivated FBS, 1X MEM NEAA and 1% pen strep. GBM cells were cultured as adherent monolayers whereas Raji cells were cultured in suspension. Cells were maintained at 37°C, in a 5% CO_2_ atmosphere and regularly sub-cultured when reaching approximately 70-80% confluence. All cell lines were routinely tested for mycoplasma contamination using a MycoAlert PLUS Mycoplasma Detection Kit (#LT07-710, Lonza, Switzerland).

### GBM cell line lentiviral transduction

Parental EGFRvIII-negative U87 and U251 cell lines were transduced with a pmp71 lentiviral vector encoding a full-length EGFRvIII, to generate stable EGFRvIII-expressing U87vIII and U251vIII cell lines, respectively. For *in vitro* cytotoxicity assays, BS153, U251 and U251vIII cells were transduced with an Incucyte NucLight Green lentivirus (#4624, Sartorius, Germany) to express a nuclear-restricted EGFP (nEGFP) fluorescence viability reporter protein. For *in vivo* tumor monitoring by bioluminescence imaging (BLi) and fluorescence labeling of tumor cells in downstream assays, U87 cells were transduced with a Luc2-mTagBFP2 lentivirus and U251vIII cells were transduced with an iRFP713-NLuc lentivirus. EGFRvIII-overexpression and fluorescence/bioluminescence vector sequences are listed in **Table S4**. For lentiviral transduction, tumor cells were seeded at 1 × 10^5^ cells per well of a 24-well plate and rested for 24 h. Growth media were replaced with antibiotic-free growth media containing 8 µg/mL polybrene. Lentiviral suspensions were added to the cells at different MOIs and incubated for 6 h at 37°C. Afterward, transduction media were replaced with fresh growth media and the cells were expanded for 1-2 weeks. Cells expressing the relevant surface receptor or fluorescence protein were sorted using a BD FACSMelody or a BD FACSAria SORP Cell Sorter (BD Biosciences, USA). Luc2 and NLuc expression was confirmed by exposing cells seeded in the wells of a flat-bottom white 96-well plate to 1 volume of 15 mg/mL D-luciferin (#LUCNA-1G, Goldbio, USA) or 1 volume of 0.5 mg/mL Nano-Glo *In Vivo* Substrate, fluorofurimazine (FFz; #CS320501, Promega, USA), respectively. Cells were imaged after a 10 min incubation protected from the light and imaged with a Fusion FX System (Vilber, France).

### Cell surface marker expression analysis

The expression of cell surface markers was determined by flow cytometry (FC). Briefly, single-cell suspensions (SCS) were counted by Trypan blue exclusion and seeded in 96-well plates at a density of 2 × 10^5^ cells per well. Cells were washed with PBS and resuspended in 100 µL of antibody staining solution. Depending on the experiment, a viability staining step was performed either before antibody staining, using Zombie NIR Viability kit (#423105, BioLegend, USA) or after, using BD Pharmingen DAPI Solution (#564907, BD Biosciences, USA) or DRAQ7 (#424001, BioLegend, USA). Antibodies, dyes and their respective dilutions are listed in **Table S5**.

### CAR T cell cytotoxicity assay

Killing assays were performed using an Incucyte S3 Live-Cell Analysis System (Sartorius, Germany). nEGFP-labeled target cells were seeded in flat-bottom, clear, 96-well plates at a density of 1 × 10^4^ cells per well and incubated for 24 h to allow the formation of a cell monolayer. mCherry-labeled CAR T cells were added at various effector-target (E:T) ratios and co-cultures were followed for 72 h. Brightfield and fluorescence images were recorded every 4 h with a 10X objective. Target cell viability kinetics were analyzed via time-lapse videos generated with Incucyte Software (Sartorius, Germany) and quantified using GraphPad Prism v10 Software (GraphPad, USA). Target cells incubated in medium alone were used to determine the baseline viability kinetics of each target cell line. Specific target cell lysis was calculated for each 4 h timepoint as green object area or count, depending on the experiment. All conditions were performed as duplicates or triplicates. Representative time-lapse videos of all assessed GBM:CAR T cell co-culture conditions are provided as **Supplementary files 1** to **10**.

### CAR T cell degranulation assay

T cell degranulation in co-cultures with GBM cells was assessed by FC. Briefly, tumor cells were washed with PBS, dissociated and counted by Trypan blue exclusion. GBM cells were seeded in a flat-bottom 96-well plate at a density of 1 × 10^4^ cells per well in 100 µL of growth medium and incubated for 24 h to allow the formation of a cell monolayer. Afterward, media in the wells was discarded, replaced by 100 µL of CAR T cell or mock-transduced T cell suspensions in GBM growth medium in 1:1 E:T and incubated for 24 h. At 24 h of co-culture, suspension cells were gently mixed in the supernatant and collected into a round-bottom 96-well plate. Cells were washed with PBS and stained with a BB700-conjugated anti-CD107a (clone H4A3; #566558, BD Biosciences, USA) for 20 min in the dark at 4°C. After surface staining, the cells were washed 3 times with autoMACS Running Buffer and resuspended in 100 µL per well of 0.5X DAPI diluted in autoMACS Running Buffer. Antibodies, dyes and their respective dilutions are listed in **Table S5**. Samples were acquired with a CytoFLEX Flow Cytometer and data were analyzed with FlowJo v10 Software. Positively stained cells were differentiated from the background using unstained controls. All conditions were performed as triplicates.

### GBM:CAR T cell co-culture supernatant ELISA

Flat-bottom F96 MAXISORP NUNC-IMMUNO plates (#439454, Thermo Scientific, USA) were coated with 50 µL per well of Purified anti-human IFNγ Antibody (clone MD-1; #507502, BioLegend, USA) diluted 1:200 in 1X PBS (#5460-0023, BioConcept, Switzerland) and incubated overnight at 4°C. The following day, wells were washed 3 times with PBS-T buffer (PBS with 0.05% Tween 20) and blocked for unspecific binding with 100 µL per well of 1% SureBlock solution (#SB232010, LubioScience, Switzerland) for 1 h at RT. After the blocking, plates were washed 3 times with PBS-T, then 50 µL of 24 h co-culture supernatants or serial dilutions of Recombinant Human IFNγ standard (#300-02, Peprotech, USA) were added to the wells and incubated for 2 h at RT. Wells were washed 3 times before adding 50 µL per well of Biotin anti-human IFN-γ Antibody (clone 4S.B3; #502504, BioLegend, USA) diluted 1:400 in 1% SureBlock solution and incubated 1 h at RT. Wells were again washed 3 times before adding 50 µL per well of HRP Streptavidin (#405210, BioLegend, USA) diluted 1:2000 in 1% SureBlock solution and incubated 1 h at RT. Wells were washed 3 times and 100 µL per well of SIGMAFAST OPD tablet (#P9187, Sigma-Aldrich, USA) solution was added. The chromogenic reactions were stopped by adding 50 µL per well of 10% sulfuric acid (H_2_SO_4_) and absorbance values were measured by a Synergy H1 Hybrid microplate reader (BioTek, USA). Absolute IFNγ concentrations in test samples were interpolated from a standard curve and represented using GraphPad Prism v10 Software.

### On-cell CD47 blocking assay

BS153 viability and count were assessed by Trypan blue exclusion, after which cells were seeded in flat-bottom 96-well plates at a density of 3 × 10^5^ cells per well. Cells were then treated for 30 min at 4°C with 50 µL per well of 10 µg/mL of InVivoMAb anti-human CD47 (clone B6.H12; #BE0019-1, Bio X Cell, USA) or InVivoMAb mouse IgG1 isotype control (clone MOPC-21; #BE0083, Bio X Cell, USA) or conditioned-media from 24 h-rested, antigen-naïve aEGFRvIII or aEGFRvIII-SGRP CAR T cells seeded at a density of 1 × 10^6^ cells per mL of unsupplemented RPMI. Pre-treated tumor cells were washed with PBS and incubated for 30 min at 4°C with biotinylated SIRPα (bt-SIRPα; #CDA-H82F2, ACROBiosystems, USA), which bound to the unblocked CD47 on BS153 cells. Finally, APC Streptavidin (SA; #405207, BioLegend, USA) staining was performed, followed by 3 washes with autoMACS Running Buffer and resuspension in 100 µL per well of 0.5X DAPI in autoMACS Running Buffer. Antibody treatments, SA stains, dyes and their respective final concentrations/dilutions are listed in **Table S5**. Samples were acquired by a CytoFLEX Flow Cytometer and data were analyzed with FlowJo v10 Software.

### Animal experimentation

All animal handling, surveillance and experimentation were performed according to the guidelines of the Swiss Federal Veterinary Office (SFVO) and the Cantonal Veterinary Office (CVO) of Basel-Stadt, Switzerland. GBM model experiments were executed under license #2929_31795 and lymphoma model experiments under license #3036_34231. Animals were maintained at the local animal facility in pathogen-free, ventilated HEPA-filtered cages under stable housing conditions of 45-65% humidity, a temperature of 21-25°C and a gradual light cycle from 7 am to 5 pm. Animals were provided standard food and water without restrictions.

### Mouse strains and humane endpoint assessment

NOD.Cg-Prkdc^scid^ Il2rg/SzJ (NSG) mice with identifier RRID:IMSR_JAX:005557 were obtained from in-house breedings or externally (Janvier Labs, France) under protocols approved by the SFVO and CVO of Basel-Stadt. Co-housed animals were assigned to treatment or control groups using a randomized approach and euthanized upon reaching the humane endpoint, including significant reduction of locomotion, significant weight loss and mild-to-severe neurologic symptoms. Scoring criteria are detailed in **Table S6**. Tumor cell implantation was set as day 0 and the survival time was set as the day of euthanasia.

### GBM mouse models and survival assessment

All experiments were performed on NSG mice of the male sex, aged 7-12 weeks at the time of tumor implantation. To assess the efficacy of anti-EGFRvIII CAR T cell and anti-CD47 antibody monotherapies, mice were injected intracranially (i.c.) with 5 × 10^4^ U251vIII-NLuc cells. To test our proposed combination therapy, mice were injected i.c. with a total of 5 × 10^4^ GBM cells consisting of 2.5 × 10^4^ U87-Luc2 and 2.5 × 10^4^ U251-NLuc, resulting in EGFRvIII-mosaic tumors. In both models, the animals were anesthetized in an induction chamber with 2.0 ± 0.5% isoflurane in an O_2_ atmosphere immediately before the tumor injections. Anesthesia on the stereotactic frame (Neurostar, Germany) was maintained at 2.0 ± 0.5% isoflurane delivered through a nose/mouth adaptor. To prevent drying of the eyes during surgery, eye gel (Lacrinorm, Bausch+Lomb Swiss AG, Switzerland) was applied. The scalp was briefly swabbed with povidone-iodide solution and a midline incision was made. A burr hole was manually drilled 2 mm lateral from the cranial midline and 1 mm posterior of the bregma suture using a microdrill. A digitally-controlled injection was performed with the Stereodrive Software (Neurostar, Germany) using a 10 µL syringe (#80300, Hamilton, USA). The syringe was lowered into the burr hole to a depth of 3 mm below the surface of the dura and retracted by 0.5 mm to form a small reservoir. A volume of 4 µL of SCS was injected at 1 µL per min. The needle was left in place for at least 1 min and carefully retracted by 0.5 mm every 30 sec. After injection, the incision was sutured (#8661H, Ethicon, USA). Buprenorphine analgesia (Bupaq-P, Streuli Tiergesundheit, Switzerland) was given intraperitoneally (i.p.) at 0.05 mg per kg immediately post-op. The following treatments and controls were administered intratumorally (i.t.) in a volume of 4 µL on days 7 and 14: vehicle (PBS), antibody isotype (InVivoMAb mouse IgG1 isotype control, clone MOPC-21; 5 µg), aCD47 (InVivoMAb anti-human CD47, clone B6.H12; 5 µg), aCD19 CAR (5 × 10^5^ cells), aCD19-SGRP CAR (5 × 10^5^ cells), aEGFRvIII CAR (5 × 10^5^ cells), aEGRFvIII CAR + aCD47 (5 × 10^5^ cells and 5 µg, respectively) or aEGFRvIII-SGRP CAR (5 × 10^5^ cells). In EGFRvIII-mosaic survival experiments, aCD47 and aEGFRvIII CAR + aCD47 treatment groups received additional doses of anti-CD47 (100 µg) administered i.p. in a volume of 100 µL on days 19, 22, 26 and 29. Antibody treatments and their respective final concentrations by application are listed in **Table S5**. Mice were monitored for clinical signs until day 90, upon which all remaining survivors were either euthanized or assigned to a tumor rechallenge experiment. Animals assigned to a rechallenge experiment were injected with the same mix of tumor cells using the same procedure described above. Tumor-rechallenged animals received no treatment and a historic vehicle group was used as a control. Kaplan–Meier survival comparison was performed using a Log-rank (Mantel-Cox) test with GraphPad Prism v10 Software.

### *In vivo* GBM bioluminescence imaging

GBM engraftment and growth were monitored by bioluminescence imaging (BLi). Mice implanted i.c. with EGFRvIII-mosaic tumors were subjected to dual BLi with specific substrates to separately monitor the growth of the EGFRvIII^+^ and EGFRvIII^-^ tumor fractions. The animals’ clinical scores and luminescence images were taken weekly, after i.p. injection of 150 mg kg^-1^ of D-luciferin or intravenous (i.v.) injection of 0.325 µmol of fluorofurimazine per mouse. Subsequently, mice were anesthetized by isoflurane inhalation and imaged on a Newton 7.0 instrument (Vilber, France). Photon flux over a defined ROI (brain surface) was quantified with NEWTON v7 Software (Vilber, France).

### Olink proteomics of mouse plasma

Peripheral blood samples were collected approximately 24 h after each treatment dose (days 8 and 15 after tumor implantation). Unanesthetized mice were briefly put under a heat lamp and then placed in a cylindrical restrainer. A small puncture was made on the tail to allow the dripping of approximately 100 µL of blood into lithium heparin-coated Microvette 100 capillary blood collection tubes (#20.1282.100, Sarstedt, Germany). Blood samples were centrifuged at 4500 rpm for 15 min at RT and the top layer of plasma was transferred into sterile 0.5 µL Eppendorf tubes. Samples were immediately frozen at -80°C until analysis. 15 samples were analyzed by proximity extension assay technology using a standard Olink Target 96 Immuno-Oncology panel (Olink Holding, Sweden). Samples were analyzed across 2 plates, 16 samples from the first run were included in the second to perform a reference sample normalization. This bridging procedure consisted in calculating the median of the paired normalized protein expression levels (NPX) differences per protein between the overlapping samples to determine the adjustment factors to be applied between the 2 datasets. For the overlapping samples, the mean value was kept. A cyclic loess normalization was applied. Since data below the limit of detection may be non-linear, the differential expression for the contrasts of interest was assessed by a Mann-Whitney U test with Benjamini-Hochberg correction. Source data is available in **Table S7**.

### Mouse tissue collection and processing

For spleen size measurement, quantification of brain tumor area, IHC and IF brain analysis, mice were anesthetized i.p. with a mix of 80 mg per kg of ketamine (Ketanarkon, Streuli Tiergesundheit, Switzerland) and 16 mg per kg of xylazine (Rompun, Elanco, USA), and transcardially perfused with ice-cold PBS. Brains and spleens were dissected and immediately fixed in formalin at 4°C for 72 h before paraffin embedding. For spectral FC analysis, mice were euthanized by CO_2_ suffocation and tumor-bearing cerebral hemispheres were harvested into ice-cold HBSS. Brain tissues were manually minced using razor blades and enzymatically dissociated at 37°C for 30 min with 1 mg/mL collagenase type IV (#LS004188, Worthington Biochemical Corporation, USA) and 250 U/mL DNase 1 (#10104159001, Roche, Switzerland) in a buffer containing HBSS with Ca^2+^/Mg^2+^ (#14205-050, Gibco, USA), 1% MEM NEAA, 1 mM sodium pyruvate, 44 mM sodium bicarbonate (#25080-060, Gibco, USA), 25 mM HEPES (#H0887, Gibco, USA), 1% GlutaMAX-I and 1% antibiotic-antimycotic (#15240062, Gibco, USA). The resulting cell suspensions were filtered through a 70 μm strainer and centrifuged in a density gradient using debris removal solution (#130-109-398, Miltenyi Biotec, Germany) according to the manufacturer’s protocol to remove myelin and cell debris. Erythrocytes were removed using ACK lysing solution (#A1049201, ThermoFisher Scientific, USA) and cell suspensions were washed with PBS and kept on ice until spectral flow staining.

### Mouse brain immunohistochemistry

#### Staining

Formalin-fixed brains were embedded in paraffin and 5 µm sections were made using a microtome. Slides were stained according to the standard Hematoxylin and eosin (H&E) protocol using the automated Gemini AS Slide Stainer (ThermoFisher Scientific, USA) and covered with Permount Mounting Medium (#SP15-100, ThermoFisher Scientific, USA). For CD3 and CD68 IHC stainings, formalin-fixed paraffin-embedded (FFPE) brain sections were stained with primary antibody in a Ventana DISCOVERY ULTRA Research Staining System (Ventana Medical Systems Inc., USA), using Histofine Simple Stain MAX PO anti-rabbit (UIP anti-rabbit; #414142F, Nicherei Biosciences Inc., Japan) as a detection reagent. Antibodies, the detection reagent and their respective final dilutions are listed in **Table S5**. Slides were counterstained with Hematoxylin II (#790-2208, Ventana Medical Systems Inc., USA) and post-counterstained with Bluing Reagent (#760-2037, Ventana Medical Systems Inc., USA). Detailed Ventana automated staining protocols for CD3 and CD68 are provided as **Supplementary files 11** and **12**.

#### Acquisition

Slides were acquired using a Nanozoomer S60 digital slide scanner (Hamamatsu, Japan) with a 40X objective.

#### Histological brain tumor size measurement

To calculate tumor size, a pixel classifier was trained to detect tumors in H&E-stained mouse brain histological sections in QuPath^39^. The Random Trees Classifier was used with the following parameters: Pixel size: 3.53 µm; Channels: red, green and blue; Features: Gaussian and Laplacian of Gaussian; Scales: 4.0 and 8.0; no normalization. Classification of tumor areas was based on example annotations to distinguish brain tissue, tumor and background (ignore*). Results were visually verified. The cumulative tumor area per sample was quantified using GraphPad Prism v10 Software. Source data are provided in **Table S8.**

#### CD3 and CD68 quantification

Cells were segmented based on Hematoxylin staining with the StarDist2D^40^ plugin within QuPath^39^ using the following parameters: Probability threshold: 0.2; Pixel size: 0.2 µm; Cell expansion: 2.0 µm; using the pre-trained model ‘he_heavy_augment.pb’. Cells were classified for CD3 and CD68 positivity based on mean UIP staining in the nucleus with a threshold of 0.12 and 0.2, respectively. All images were prepared using OMERO.web app (www.openmicroscopy.org/omero/figure/). The tumor core was delineated based on nuclear stain density and the tumor rim was defined as a 100 μm-wide area surrounding the tumor core. Relative cell positivity per brain section was calculated as a percentage of stained vs non-stained cells in areas defined as tumor core or tumor rim and quantified using GraphPad Prism v10 Software. Source data are provided in **Table S8.**

### Mouse brain immunofluorescence multiplexing

#### Staining

The protocol was adapted from Gut and colleagues^41^. FFPE brains were sectioned in 5 µm-thick sections using a microtome. Slides were deparaffinized 3 times for 5 min with ROTI Histol (#6640, Carl Roth, Germany) and 30 sec with 100% EtOH, 95% EtOH, 70% EtOH, 50% EtOH and dH_2_O washing steps. For antigen retrieval, slides were exposed to pre-heated 1X Citrate Buffer (#C9999, Sigma-Aldrich, USA), followed by a washing step with PBS for 10 min. Tissue sections were permeabilized in PBS with 0.2% Tween-20 (#P5927, Sigma-Aldrich, USA) twice for 15 min. Thereafter, the slides were washed with PBS twice for 10 min followed by blocking for 1 h in a dark humid chamber with 10% normal donkey serum (#017-000-121, Jackson ImmunoResearch, USA) diluted in PBS. Afterward, the slides were stained in different cycles overnight at 4°C with the following antibodies: anti-CD3 (clone CD3-12; #MCA1477, Bio-Rad, USA), anti-CD206 (clone E6T5J; #24595S, Cell Signaling Technology, USA), anti-EGFRvIII (clone RM419; #MA5-36216, ThermoFisher Scientific, USA), anti-GFAP (clone D1F4Q; #12389S, Cell Signaling Technology, USA), anti-IBA1 (polyclonal; #NB100-1028, Novus Biologicals, USA), anti-Ki67 (clone SolA15; #14-5698-82, ThermoFisher Scientific, USA) and anti-TMEM119 (clone 28-3; #ab209064, Abcam, UK). After each primary antibody staining cycle, the slides were washed 3 times for 5 min with PBS and then stained with the respective secondary fluorescent antibodies and DAPI for 1 h in a dark humid chamber at RT. Antibodies, dyes and their respective final dilutions are listed in **Table S5**. The slides were washed 3 times for 5 min with PBS and mounted with imaging buffer at pH 7.4 (700 mM N-Acetyl-Cysteine (NAC; #160280250, ThermoFisher Scientific, USA) in ddH_2_O + 20% HEPES solution (#H0887, Sigma-Aldrich, USA)). For antibody elution after each imaging cycle, the slides were treated 3 times for 15 min with elution buffer containing 0.5 M L-Glycine (#3790.2, Carl Roth, Germany), 1.2 M Urea (#U5378, Sigma-Aldrich, USA), 3 M Guanidium chloride (#G3272, Sigma-Aldrich, USA), and freshly-added 70 mM TCEP-HCl (#C4706, Sigma-Aldrich, USA), followed by a ddH_2_O wash and restaining.

#### Acquisition

Images were acquired using an ECLIPSE Ni upright microscope (Nikon, Japan) equipped with 395, 470, 561 and 640 nm lasers with a Prior PL-200 robotic slide loader equipped with Microscan MiniHawk (Omron Microscan Systems Inc., Japan) and the Photometrics Prior 95B camera (Teledyne Photometrics, USA). The acquisition was done using JOBS automation in NIS-Elements v5.11.00 Software. 4X Plan Apo NA 0.2 (Nikon, Japan) objective was used to make a slide overview and ‘general analysis3’ with ‘Otsu’ threshold was used to detect the tissue. The focus surface was created by performing software autofocus every 2 points based on the DAPI signal with Plan Apo λ 20X NA 0.8 and the tissue was scanned with the same objective. Slides were scanned after each staining cycle, with blank (unstained) cycles used to evaluate autofluorescence. The exposure time of each channel was constant for all cycles and corresponding blank channels (GFP, Cy3 and Cy5) were subtracted from the signal cycle.

#### Image preprocessing

Fiji^42^ was used for image preprocessing. Single tiles were stitched using Grid/Collection Stitching^43^. Stitched tissues were registered based on the DAPI channel in consecutive cycles using MultiStackReg plugin^44^ and the same correction was propagated on the remaining channels. Images were saved as a pyramidal file using the Kheops^45^ plugin, available under an open-source license: https://github.com/BIOP/ijp-kheops/releases.

#### Cell segmentation and quantification

Cells were segmented based on DAPI staining with the StarDist2D^40^ plugin within QuPath^39^ using the following parameters: Probability threshold: 0.5; Pixel size: 0.5 µm; Cell expansion: 2.0 µm; using a pre-trained model ‘dsb2018_heavy_augment.pb’. Object classification in QuPath was used to define cell positivity for CD3, CD206, EGFRvIII and IBA1. All images were prepared using OMERO.web app (www.openmicroscopy.org/omero/figure/). Cell population ratios per treatment condition were quantified using GraphPad Prism v10 Software. Source data are provided in **Table S8.**

### Spectral flow cytometry

#### Staining and acquisition

After the ACK-lysis step described in the “Mouse tissue collection and processing” section, freshly dissociated cells were washed with PBS and resuspended in 400 µL of PBS. A 200 µL fraction of one sample per treatment group was used as an unstained control to detect and correct for condition-specific autofluorescence. In all cases, input volume for full-stained samples were kept at 200 µL to ensure equivalent staining conditions across all samples. The remaining 200 µL of SCS volume was mixed and used to generate fluorescence minus one (FMO) stainings, allowing for condition-specific FMO gating. All centrifugation and incubation steps were performed at 300 × g at 4°C, protected from light, unless otherwise stated. Viability staining was performed by incubating cells with Zombie NIR Viability kit for 20 min. Subsequently, Fc-block was performed by incubating cells in a dilution of Human TruStain FcX (#422302, BioLegend, USA) and mouse TruStain FcX (#101320, BioLegend, USA) for 10 min. Antibody mastermixes (full-stains and FMOs) were freshly prepared on the day of staining in Brilliant Stain Buffer (#00-4409-75, ThermoFisher Scientific, USA). Cell surface markers were stained with surface antibody mastermixes for 25 min and then washed twice in autoMACS Running Buffer. To allow the subsequent staining of intracellular antigens, a fixation/permeabilization step was performed by incubating the cells for 20 min at RT using a Cyto-Fast Fix/Perm Buffer set (#426803, BioLegend, USA). Intracellular antibody staining was then performed for 25 min at RT. The spectral FC staining panel is summarized in **Table S5.** Finally, the samples were washed twice, resuspended in a final volume of 200 µL of PBS and acquired on a Cytek Aurora 5-Laser Spectral Analyzer (Cytek Biosciences, USA) using standard, daily quality-controled, Cytek-Assay-Settings.

#### Data preprocessing

Spectral unmixing was performed using SpectroFlo v3.1.0 Software. UltraComp eBeads Plus Compensation Beads (#01-3333-42, ThermoFisher Scientific, USA) were used as single reference controls (SRC). SRC for mCherry and mTagBFP2 were generated with mock-transduced and the respective transduced, CAR T cells (mCherry) and U87 tumor cells (mTagBFP2) from cell cultures. Due to the fluorescent nature of our unstained single cell suspensions (mCherry^+^ CAR T cells and mTagBFP2^+^ tumor cells) and to avoid contamination of the autofluorescence signature of all immune populations, fcs files of ‘unstained’ conditions were first cleaned by exclusion of V3^hi^ mTagBFP2 tumor cells and YG3^hi^ mCherry CAR T cells. The resulting fcs files were then reimported and used to detect populations with highly divergent autofluorescence signatures by NxN plots. When detected, such populations were extracted in the same manner and separately added as a fluorescence tag to avoid assigning incorrect marker expression caused by heterogenous autofluorescence between immune cell populations. Conventional analysis and generation of input fcs files (gates: debris removal, single cells, live cells and CD45^+^ cells) for downstream analysis with R was performed using FlowJo v10 Software and is detailed in the “FlowSOM analysis” section below.

#### FlowSOM analysis

Data were manually pre-gated to remove the debris and select for CD45^+^, live, single cells using FlowJo v10 Software. The analysis was subsequently performed in R (version 4.3.1). Data was transformed by asinh transformation using variance stabilizing cofactors for each channel (*estParamFlowVS* and *transFlowVS* functions from the FlowVS package), except for the mTagBFP2 channel where the cofactor was manually set to 3. Pre-processing QC (using *PeacoQC* function from the PeacoQC package) was performed to remove outliers and unstable events (IT_limit was set at 0.55 and MAD at 6). Clustering was performed using FlowSOM and ConsensusClusterPlus using the wrapper function *cluster* from the CATALYS package (xdim = 10, ydim = 10, maxK = 15, seed = 1234). The resulting clusters were manually annotated. Differential testing was performed using the diffcyt package (diffcyt-DA-edgeR for differential abundance and diffcyt-DS-limma for differential state).

#### Conventional flow cytometry data analysis

FlowJo v10 Software was used for data analysis. Gates were set using FMO controls. Either the percentage of cell population of interest or median fluorescence intensity (MFI) was reported.

#### Human GBM samples and patient information

Freshly resected primary tumor tissue samples from patients with a pathology-confirmed GBM diagnosis were obtained from the Neurosurgical Clinic of the University Hospital Basel, Switzerland, following the Swiss Human Research Act and institutional ethics commission (EKNZ 02019-02358). The study was conducted following the ethical principles of the Declaration of Helsinki, regulatory requirements, and the Code of Good Clinical Practice. All patients gave written informed consent for tumor biopsy collection and signed a declaration enabling the use of their biopsy specimens in scientific research (Req-2019-00553) with all identifying information removed. All patients were treatment-naïve at the time of tissue collection. The clinical information of GBM patients is summarized in **Table S9**.

#### Human GBM tissue processing

GBM tissue samples were transported on ice and transferred to the laboratory for dissociation into single-cell suspensions within 2-3 h after surgical resection. Human brain tissue was mechanically minced using razor blades and enzymatically dissociated as described under “Mouse tissue collection and processing”, except that debris and myelin were removed by a 0.9 M sucrose (#84100, Sigma-Aldrich, USA) density gradient centrifugation. After ACK-lysis, the single-cell suspensions were washed with PBS, resuspended in Bambanker at an approximate density of 2 × 10^6^ live cells per mL and stored long-term in LN_2_.

#### Real-time quantitative PCR

The total RNA contents of approximately 2 × 10^6^ cells from culture-dissociated tumor cells or human GBM single-cell suspensions were extracted with TRIzol Reagent (#15596026, Invitrogen, USA) and a Direct-zol RNA Miniprep Plus kit (#R2070, Zymo Research, USA). cDNA was synthesized with a SuperScript VILO cDNA Synthesis Kit (#11754-050, ThermoFisher Scientific, USA) using 20 ng of input RNA in a 20 µL reaction. The reaction included priming for 10 min at 25°C, reverse transcription for 60 min at 42°C and inactivation for 5 min at 85°C. RT-qPCR reactions were performed with a SsoFast Eva Green supermix (#172-5201, Bio-Rad, USA) using 2 µL of input cDNA in a 20 µL reaction. RT-qPCR primers are listed in **Table S10**. Thermal cycling was performed using a CFX96 Touch Real-Time PCR Detection System (Bio-Rad) and included an initial step of enzyme activation for 30 sec at 95°C, followed by 40 cycles of denaturation for 5 sec at 95°C and annealing/extension for 5 sec at 60°C. Relative EGFR and EGFRvIII expression was calculated with the ΔΔCt method, using GAPDH Ct to normalize signal expression.

#### Pharmacoscopy

Frozen patient single-cell suspensions were thawed and resuspended in (DMEM #11995065, Gibco, USA) supplemented with 10% FBS, 25 mM HEPES and 1% pen strep and were seeded at 8 × 10^3^ cells per well in 50 µL per well into clear-bottom, tissue-culture treated, CellCarrier-384 Ultra Microplates (#50-209-8071, Perkin Elmer, USA). mCherry-labeled CAR T cells were added and co-cultured with the patient single cell suspensions for 48 h at 37°C, 5% CO_2_. Every CAR T cell co-incubation condition was tested with five technical replicate wells and six technical replicate wells for the PBS control. After the co-incubation period, the cells were fixed with 4% PFA, blocked with PBS containing 5% FBS, 0.1% Triton-X and 4 µg/mL DAPI (#422801, BioLegend, USA) for 1 h at RT and stained with two staining panels: (1) anti-CD3 (clone UCHT1; #300415, BioLegend, USA) and anti-CD14 (clone HCD14; #325612, BioLegend, USA); and (2) anti-NESTIN (clone 10C2; #656802, BioLegend, USA) and anti-EGFRvIII (clone RM419; #MA5-36216, ThermoFisher Scientific, USA) overnight at 4°C. The wells were washed, and the cells that were stained with primary antibodies were incubated with the respective secondary fluorescent antibodies for 1 h at RT in the dark. The staining panels are summarized in **Table S5**. The plates were imaged with an Opera Phenix automated spinning-disk confocal microscope at 20X magnification (Perkin Elmer, USA). Single cells were segmented based on nuclear DAPI staining with CellProfiler v2.2.0 Software. Downstream image analysis was performed with MATLAB R2021b. Marker-positive cell counts for each condition were identified based on a linear threshold of each channel, were averaged across each well and compared between CAR T cell co-incubation conditions. Source data are available in **Table S11**.

#### Lymphoma mouse model and survival assessment

To test the efficacy of the CAR constructs in a subcutaneous (s.c.) tumor model, NSG mice of the female sex, aged 8-12 weeks were anesthetized with isoflurane and injected with 5 × 10^5^ CD19-expressing Raji cells s.c. in the right flank. Raji cells were suspended in BD Matrigel Basement Membrane Matrix High Concentration (#354248, BD Biosciences, USA) diluted 1:1 in phenol red-free DMEM (#31053028, Gibco, USA) without additives in a total volume of 100 µL. Three days after tumor inoculation, each mouse received 8 × 10^5^ CAR T cells suspended in 200 µL of PBS or PBS alone, i.v. via the tail vein. Tumor-bearing mice injected with unspecific CAR T cells or PBS were used as controls. Tumor size was measured 3 times per week using a caliper. Animals were sacrificed before reaching a tumor volume of 1500 mm^3^ or when reaching an exclusion criterion (ulceration, severe weight loss, severe infection or bite wounds). Tumor volume was calculated according to the following formula: Tumor volume (mm^3^) = (d^2^ × D)/2 with D and d being the longest and shortest tumor parameter in mm, respectively.

### Statistical analyses

Data analysis and visualization were performed using Excel version 16.14.1 (Microsoft) and Prism 8.4 (GraphPad). Graphs represent either group mean values ± s.d. (for *in vitro* experiments) or ± s.e.m. (for *in vivo* experiments) or individual values. For *in vitro* studies, statistical comparisons were made with either unpaired *t*-tests when comparing two groups or one-way ANOVA with multiple comparison corrections when comparing more than two groups. For *in vivo* studies, survival curves were compared with the log-rank test; tumor growth was compared with repeated-measures ANOVA and the Mann–Whitney test was used to compare two groups. *P* < 0.05 was considered statistically significant. *P* values are denoted with asterisks: * *P* < 0.05; ** *P* < 0.01; *** *P* < 0.001; and **** *P* < 0.0001.

## Results

### Engineering CAR T cells that secrete a SIRPγ-derived blocker of the CD47-SIRPα phagocytosis axis

Antigen escape is a major mechanism of anti-EGFRvIII CAR T cell therapy resistance (**Fig. 1A**). We propose a fourth-generation CAR design, whereby anti-EGFRvIII CAR T cells constitutively release a soluble SIRPγ-related protein (SGRP) with high affinity to CD47^30^ (**Fig. 1B**). To design a CAR T cell that concomitantly targets GBM cells via EGFRvIII recognition and reprograms GAMs by disrupting the CD47-SIRPα phagocytosis axis, we equipped anti-EGFRvIII-BBz CAR T cells (3C10.BBz^34^, termed aEGFRvIII CAR) with a secretable SGRP by adding an IL2 signal peptide to the SGRP sequence, resulting in the generation of aEGFRvIII-SGRP CAR (**Fig. 1C**). Synthetic SGRP shares a strong identity with the endogenous SIRPγ binding domain (SIRPγ-V1) amino acid (AA) sequence (**Fig. S1A** and **Table S1**) and binds to human CD47 in a similar manner as that of competitive human and murine SIRPα (**Fig. S1B**). Specifically, we generated *EF1A*-driven polycistronic constructs encoding overall identical CAR structures targeting either EGFRvIII or control antigen CD19 (FMC63.BBz^36^; termed aCD19 CAR), a mCherry (mC) fluorescent reporter, and a secretable SGRP. The resulting polyproteins were generated by flanking T2A or P2A self-cleaving peptide sequences (**Fig. 1D** and **Table S2**) and were incorporated into lentiviral expression vectors (**Fig. S1C**).

Activated T cells were stably transduced with lentiviral supernatants and T cell cultures were expanded in the presence of IL-2 for 5 to 7 days, subsequently enriched for mCherry-expression by cell sorting, and further expanded for downstream applications (**Fig. 1E**, **Fig. S1D** and **Fig. S1E**). As intended, aCD19 and aEGFRvIII CARs showed specific binding affinity to their respective target, but not to mismatched target proteins (**Fig. 1F** and **Fig. 1G**) or wild type (wt) EGFR protein (**Fig. S1F**).

We then tested whether SGRP was constitutively expressed by SGRP-producing CAR T cells. In conditioned media from CAR T cell cultures from 2 donors, SGRP was the most differentially enriched protein in the secretome of aEGFRvIII-SGRP CAR compared to aEGFRvIII CAR (**Fig. 1H** and **Fig. S1G**). A full list of detected proteins and their relative abundance is shown in **Table S3**. The identified peptide sequences included SGRP-specific AA modifications, confirming specific SGRP detection rather than contamination with peptides of highly conserved SIRP-family proteins (**Fig. S1H**).

### CAR T cell effector function against GBM is unaffected by SGRP-mediated CD47 blockade *in vitro*

To identify GBM cell line targets for *in vitro* CAR T cell efficacy validation, we profiled surface expression of our targets of interest in four GBM cell lines (U251, U87, U251vIII with transgenic (tg) overexpression of EGFRvIII, BS153 with endogenous EGFRvIII expression), one Burkitt’s lymphoma cell line (Raji) and one normal neural stem cell (NSC) line (NSC197). We used aCD19 CAR and aCD19-SGRP CAR as controls, and Raji as CD19^+^ target cells throughout the study (**Fig. S2A**, left plot). We confirmed the surface expression of EGFRvIII on U251vIII and BS153 (99.3% and 28.9%, respectively), while U251 and U87 showed substantially lower expression levels (16.4% and 23.5%, respectively; **Fig. S2A**, center plot). Before any *in vitro* and *in vivo* experiments, U251vIII and BS153 were sorted for their EGFRvIII^+^ cell population, while U251 and U87 were sorted for their EGFRvIII^-^ cell fraction (**Fig. S2B**). All cell lines were validated for EGFRvIII expression throughout the study. Furthermore, we also confirmed CD47 overexpression in all tumor cell lines (U251vIII: 56.5%, U251: 51.4%, U87: 53.5%, BS153: 80.6% and Raji: 58.4%), compared to NSCs used as a normal brain cell control (**Fig. S2A**, right plot).

We then established co-cultures of mCherry^+^ conventional CAR T or SGRP-secreting CAR T cells with GBM cells expressing a nuclear-restricted EGFP (nEGFP) as a fluorescence viability reporter. A decline of the nEGFP signal in the co-cultures was an indicator of tumor viability reduction, as previously validated by puromycin selection (**Fig. S2C**). Time-lapse imaging revealed the specific cytolytic effect of aEGFRvIII CAR and aEGFRvIII-SGRP CAR against U251vIII cells (**Fig. 2A**). aEGFRvIII CAR also showed similar efficacy against BS153 cells (**Fig. S2D**; representative co-culture time-lapse videos provided as **Supplementary files 1** and **2**). By contrast, no measurable off-target cytotoxicity was observed against U251, an EGFRvIII-negative-sorted GBM cell line (**Fig. 2B**). At various time points and effector-target (E:T) ratios, only target-specific (aEGFRvIII +/- SGRP) CAR T cells displayed a dose-dependent effect against EGFRvIII^tg^ GBM cells (**Fig. 2C**). Moreover, the CAR T cell cytolytic capacity remained unaffected by SGRP secretion in co-cultures with U251vIII (**Fig. 2C**; time-lapse videos provided as **Supplementary files 3** to **6**) or U251 (**Fig. S2E**; time-lapse videos provided as **Supplementary files 7** to **10**), as shown by the analogous response of aEGFRvIII CAR and aEGFRvIII-SGRP CAR in co-cultures with GBM cells in the absence of tumor-associated myeloid cells. Notably, our CAR T cell production and expansion protocol yielded almost exclusively CD4^+^ T cells (**Fig. S2F**).

**Figure 2.**
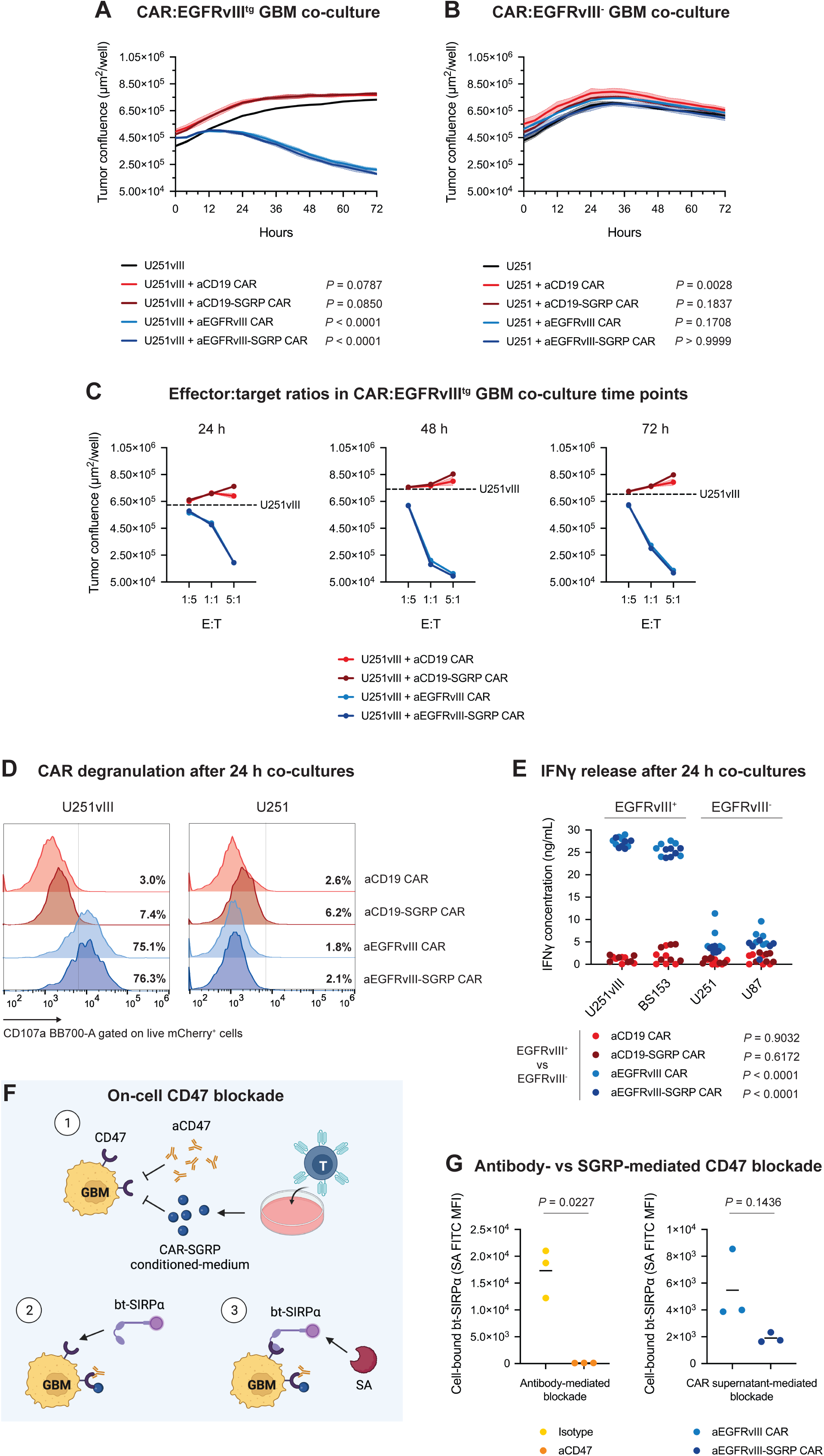
Anti-EGFRvIII-SGRP CAR T cells have comparable *in vitro* efficacy and on-target activation to conventional anti-EGFRvIII CAR T cells. **A,** Assessment of CAR T cell on-target killing capacity by co-culture time-lapse of nEGFP^+^ U251vIII with mCherry^+^ target-specific (aEGFRvIII CAR +/- SGRP) or nonspecific (aCD19 CAR +/- SGRP) at a 1:1 E:T ratio for 72 h. **B,** Assessment of CAR T cell off-target killing by co-culture time-lapse of nEGFP^+^ U251 with mCherry^+^ CAR T cells at a 1:1 E:T ratio for 72 h; **A,** and **B,** Differences between each co-culture compared to the ‘tumor alone’ control condition were analyzed using one-way ANOVA and Dunnett’s multiple comparisons tests. **C,** Dose-dependent CAR T cell killing capacity in a co-culture with U251vIII at defined time points. Dashed lines represent the mean confluence in control wells with only U251vIII cells; **A,** to **C,** Curves and data points represent the mean of duplicate measurements. Curve-adjacent shaded areas represent standard deviation. **D,** Representative histograms displaying the percentage of CAR T cell degranulation (CD107a expression) in co-cultures with U251vIII, BS153, U251 or U87 GBM cell lines for 24 h (gated on live mCherry^+^ singlets). Conditions were performed in duplicates with n = 2 HDs and the experiment was repeated once. **E,** IFNγ release detected by ELISA in supernatants of CAR T cells co-cultured with GBM cell lines expressing endogenous EGFRvIII (BS153), overexpressing EGFRvIII (U251vIII) or EGFRvIII^-^ (U251 and U87). Differences in IFNγ release between EGFRvIII^+^ and EGFRvIII^-^ co-cultures with each CAR were determined using Two-way ANOVA and Šidák’s multiple comparisons tests. Conditions were performed in triplicates with n = 2 HDs and the experiment was repeated once. **F,** Schematic illustration of the experimental setup of an SGRP/aCD47 blocking assay on CD47^+^ target tumor cells where (1) BS153 were treated with aEGFRvIII-SGRP CAR conditioned-medium or aCD47, (2) exposed to bt-SIRPα which competitively bound to available CD47, and (3) CD47-bound bt-SIRPα was assessed by SA-FITC and MFIs calculated. **G,** Representative dot plots depicting the CD47-blocking capacity of aCD47 or SGRP determined by FC-based detection of SA-FITC coupled to GBM-bound bt-SIRPα. Conditions were performed in triplicates and the experiment was repeated once.

We further examined the degranulating and cytokine-producing capacity of CAR T cells in co-cultures with GBM cell lines of varying EGFRvIII status. Flow cytometry (FC) analysis of CAR T cells in 24 h co-cultures showed that target-specific CAR T cells expressed higher levels of CD107a in response to EGFRvIII^+^ GBM cell lines (**Fig. 2D**). We also detected significantly higher concentrations of IFNγ in co-cultures of target-specific CAR T cells with EGFRvIII^+^, but not with EGFRvIII^-^ GBM cell lines (**Fig. 2E**).

To determine the CD47-blocking capacity of T cell-secreted SGRP compared to anti-CD47 antibody (aCD47) *in vitro*, we pre-treated BS153 tumor cells with CAR T cell-conditioned media (aEGFRvIII-SGRP CAR vs aEGFRvIII CAR), or antibody dilutions (anti-CD47, clone B6.H12 vs. IgG1 isotype, clone MOPC-21). In a second step, BS153 cells were exposed to biotinylated SIRPα (bt-SIRPα) that bound to the remaining available CD47 on tumor cells. Cell-bound bt-SIRPα was detected by labeling with fluorescence-conjugated streptavidin (SA). A schematic illustration of the experiment is shown in **Fig. 2F**. aCD47 treatment induced a potent blockade of the CD47-SIRPα interaction (**Fig. 2G**, left dot plot) whereas treatment with aEGFRvIII-SGRP CAR-conditioned medium slightly impaired this interaction (**Fig. 2G**, right dot plot). Since CD47 is ubiquitously expressed^21^, we hypothesize that SGRP may be partially captured in autocrine or paracrine loops by CAR T cells. Consequently, the amount of available SGRP collected from cell culture supernatants may be reduced, contributing to the apparent lower efficacy of SGRP-mediated CD47 blockade. Nevertheless, this artificial *in vitro* model overlooks crucial cell-cell interactions in the context of the GBM TME. Thus, understanding the full effect of aEGFRvIII-SGRP CARs in a wider translational context requires an orthotopic model of EGFRvIII-heterogenous GBM with GAM recruitment and local CAR T cell delivery.

### Anti-EGFRvIII-SGRP CAR therapy leads to significant survival benefit in an aggressive EGFRvIII-mosaic GBM xenograft model

Many aEGFRvIII CAR preclinical studies show effective cytotoxicity but fail to model the heterogeneity of human GBM^46,47^. Although preclinical aEGFRvIII CAR monotherapy has demonstrated efficacy, this strategy translates poorly in the clinical setting^10^. To establish a baseline for our proposed combination therapy, we first tested the efficacy of aEGFRvIII CAR and aCD47 monotherapies in an EGFRvIII^+^ GBM xenograft model. For these experiments, U251vIII cells were orthotopically implanted in NSG mice (**Fig. S3A** and **Fig. S3B**). The therapeutic scheme included i.t. treatments of mouse IgG1 (MOPC-21; isotype), anti-human CD47 (B6.H12; aCD47), aCD19 CAR and aEGFRvIII CAR (**Fig. S3B**). Tumor-bearing mice received 2 identical intratumoral (i.t.) doses of treatment or control, on days 7 and 14 post tumor implantation. Clinical scores and tumor growth were monitored weekly by bioluminescence imaging until a maximum of 13 weeks (90 days) or until animals reached the humane endpoint (**Fig. S3C**). Overall survival analysis (**Fig. S3D**) showed that local aCD47 monotherapy failed to improve survival (median survival: 34.5 days) when compared to the isotype antibody control group (median survival: 30.5 days). Conversely, aEGFRvIII CAR treatment (median survival: 65.5 days) led to 40% survival by day 90 after tumor implantation, a stark contrast to aCD19 CAR control (median survival: 29 days). Bioluminescence assessment confirmed an overall lower tumor burden in aEGFRvIII CAR-treated mice (**Fig. S3E**). For BLi-based tumor growth monitoring, the same head region of interest (ROI) was applied to every animal throughout the study (**Fig. S3F**).

To assess the efficacy of CAR + SGRP combination therapy against GBM, we established a preclinical EGFRvIII-mosaic GBM model with orthotopic tumor implantation and intracranial therapy administration in mice. First, we sorted GBM cell lines for their EGFRvIII^+^ (BS153 and U251vIII) or EGFRvIII^-^ (U87 and U251) cell fractions based on cell surface staining and then assessed their engraftment in NSG mice. We selected U251vIII and U87 for their consistent engraftment and fast progression to the onset of clinical signs. Thereafter, we co-injected U251vIII and U87 intracranially in a 1:1 ratio as single-cell suspensions and allowed them to settle for 7 days. All animals received 2 identical i.t. doses of treatment or control, on days 7 and 14 post tumor implantation. For better equipoise in comparing antibody-based CD47 blockade (aCD47) with constitutive CAR-mediated SGRP release, animals receiving aCD47 as monotherapy or in combination with aEGFRvIII CARs (aEGFRvIII CAR + aCD47) received 4 additional doses of antibody delivered i.p. within the 2 weeks following the i.t. treatment regimen. To monitor U251vIII and U87 cell populations separately *in vivo*, we differentially labeled U251vIII with NanoLuciferase (NLuc) and U87 with Luciferase2 (Luc2) bioluminescence reporters using lentiviral transgene delivery (**Table S4**). The experiment timeline and therapeutic setup are illustrated in **Fig. 3A** and **Fig. 3B**, respectively. Overall survival analysis (**Fig. 3C**) showed that untreated animals in the vehicle control group had a median survival of 31.5 days. Surprisingly, combined locoregional/systemic aCD47 monotherapy failed to improve survival (median survival: 34 days) when compared to an isotype antibody control group (median survival: 32 days). We reason that despite combined local/systemic administration, the aCD47 treatment regimen did not maintain a sufficient blockade of CD47 for effectively inducing persistent GAM modulation. Notably, aggressive tumor models in the NSG context are difficult to treat using even higher doses of aCD47 antibodies^23^. Predictably, aCD19 CAR treatment resulted in poor outcomes (median survival: 36 days). Similarly, survival after aCD19-SGRP CAR therapy (median survival: 40 days) suggested that peripherally released SGRP without direct CAR-mediated targeting was insufficient to eliminate orthotopic xenograft tumors. Nonetheless, aEGFRvIII CAR monotherapy was able to eliminate GBM in 20% of treated animals (**Fig. 3C** and **Fig. 3D**). However, the combination of aEGFRvIII CAR + aCD47 failed to improve survival compared to aEGFRvIII CAR monotherapy (20%). This pointed towards a potential elimination of grafted CD47^+^ aEGFRvIII CAR T cells via phagocytosis or insufficient continuous dosing of local anti-CD47 antibodies. Notably, clone B6.H12 was used in the experiments because of the limited availability of the superior anti-CD47 clone Hu5F9-G4^48^. In contrast, aEGFRvIII-SGRP CAR was extraordinarily potent in this challenging GBM model. A near-complete therapeutic response was observed with 94.7% overall and 63.2% tumor-free survival (**Fig. 3C** and **Fig. 3D**, respectively). Furthermore, differential bioluminescence monitoring of either U251vIII-NLuc (**Fig. 3E**) or U87-Luc2 (**Fig. 3F**) confirmed that a majority of EGFRvIII^-^ tumors were cleared after aEGFRvIII-SGRP CAR treatment. Bioluminescence plots of individual treatment groups are shown in **Fig. S4A**. A direct bioluminescence imaging comparison and quantification of EGFRvIII^+^ and EGFRvIII^-^ tumor burden in aEGFRvIII CAR, aEGFRvIII CAR + aCD47 and aEGFRvIII-SGRP CAR treatment groups are shown in **Fig. S4B** and **Fig. S4C** on a representative time point (week 7 post tumor implantation). Rechallenging of cured mice with EGFRvIII-mosaic tumors resulted in prolonged survival compared to the historic vehicle control group, pointing towards a potential persistence of aEGFRvIII-SGRP CAR T cells and/or anti-tumor GAMs in the brains of these animals (**Fig. 3G**). Interestingly, tumor monitoring after rechallenge revealed preferential control of EGFRvIII^+^ tumor growth and outgrowth of EGFRvIII^-^ clones (**Fig. S4D**). Altogether, these results demonstrate that constitutive CAR-secreted SGRP is a superior combination partner to EGFRvIII-specific CAR T cells compared to antibodies delivered locally. Owing to the ubiquitous expression pattern of CD47, the presence of competing host anti-CD47-consumer cells (e.g. neurons, glia and brain-infiltrating immune cells) probably contributes to the significant difference in efficacy observed for CAR + aCD47 or CAR + SGRP combination therapies. The dosage and persistence of CD47 blockade are equally important factors for this outcome. During antigen-positive tumor cell lysis, aEGFRvIII-SGRP CAR T cells come into close contact with antigen-negative bystander tumor cells. We hypothesize that the extended secretion of SGRP in these tumor niches during tumor cytolysis is a crucial factor in the efficacy of the aEGFRvIII-SGRP CAR treatment modality in GBM-mosaic xenografts.

**Figure 3.**
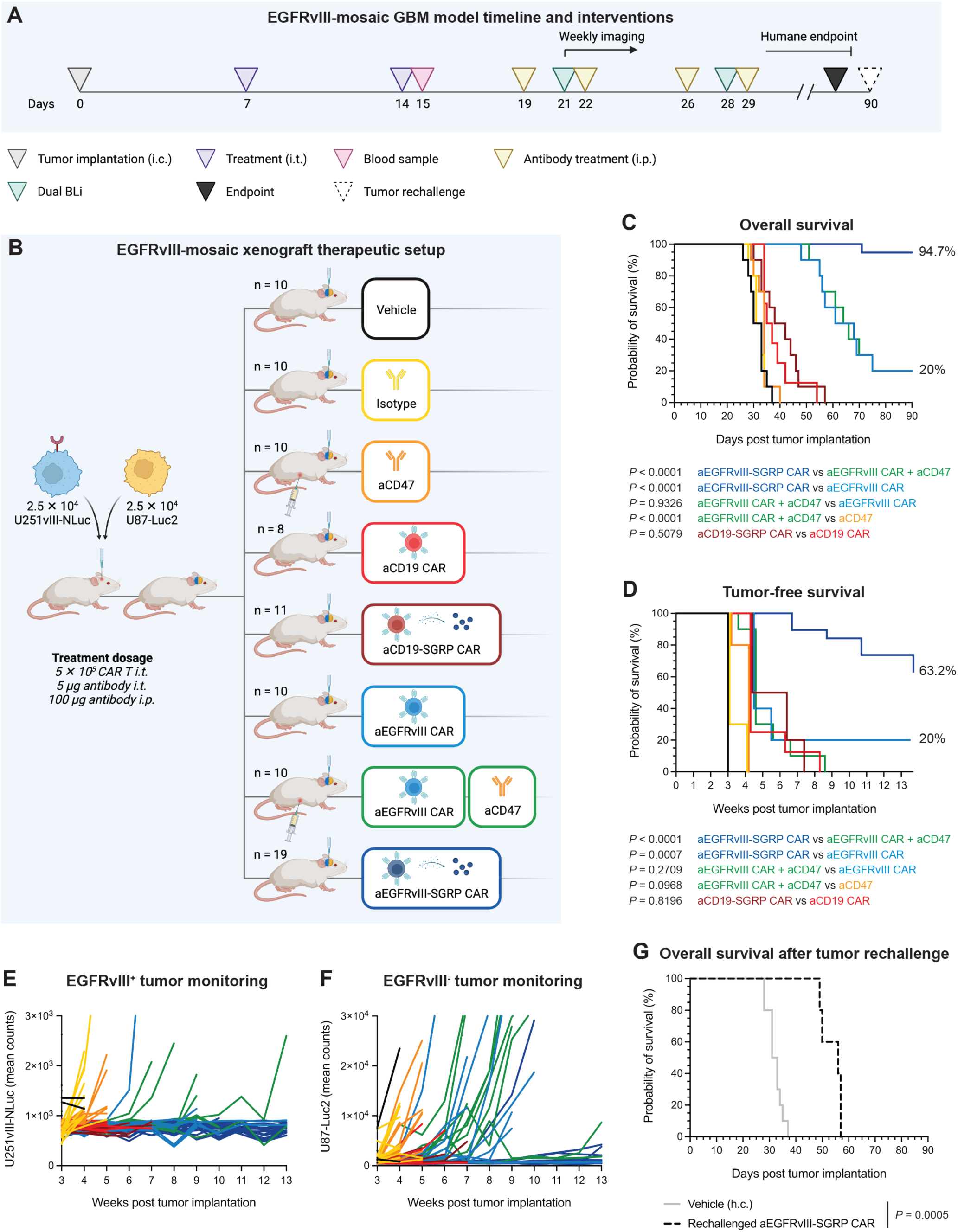
Local anti-EGFRvIII-SGRP CAR T cell therapy leads to survival benefit and long-term tumor control in an EGFRvIII-heterogenous GBM xenograft model. **A,** Experimental treatment and monitoring schedule. Animals were treated twice – at 7 and 14 days – after intracranial tumor implantation using the same stereotactic coordinates, and routinely monitored for clinical signs, weekly dual bioluminescence imaging (BLi) and morbidity/survival assessment. Plasma for cytokine analysis was collected on day 15 – 24 h after the second intratumoral (i.t.) treatment. Anti-CD47 therapy was prolonged for 4 additional intraperitoneal (i.p.) injections on days 19, 22, 26 and 29. Animals were euthanized upon reaching the humane endpoint. Upon reaching 90 days of tumor-free survival, 5 aEGFRvIII-SGRP CAR-treated animals were tumor-rechallenged in the contralateral hemisphere using the same stereotactic coordinates. **B,** Experimental setup of orthotopic xenograft experiments in NSG mice encompassing co-implanted EGFRvIII^+^ U251vIII and EGFRvIII^-^ U87 GBM cell lines mimicking tumor heterogeneity, and therapeutic/control cohorts including local CAR T cell or antibody monotherapies or combinations with local SGRP or local and systemic CD47 blockade. **C,** Kaplan–Meier plot of overall survival (in days). **D,** Kaplan–Meier plot of tumor-free survival (in weeks), combining survival assessment with BLi monitoring scores; **C,** and **D,** Log-rank tests were used to compare the indicated treatment/control groups. **E,** and **F,** Tumor progression in EGFRvIII-mosaic xenografts was monitored for each mouse using differential BLi time course imaging with either FFz or D-luciferin substrates (in weeks); **E,** U251vIII NLuc-reporter BLi curves; **F,** U87 Luc2-reporter BLi curves; **C,** to **F,** Data were pooled from 3 independent experiments. **G,** Kaplan–Meier plot of overall survival of aEGFRvIII-SGRP CAR-cured, tumor-rechallenged animals (in days). A Log-rank test compared the rechallenge group to the historic vehicle control group (vehicle (h.c.)).

### Anti-EGFRvIII-SGRP CAR therapy induces a peripherally-detected inflammatory response in GBM-bearing mice

To elucidate potential mechanisms of innate immune activation upon locoregional CAR T cell GBM treatments, we conducted an immuno-oncology-targeted proteomic analysis of plasma samples collected 24 h after the second CAR treatment (day 15 post tumor implantation). Using an Olink Immuno-Oncology panel, we analyzed a total of 92 proteins on 2 complementary datasets encompassing 54 plasma samples from individual mice (**Fig. S5A**). Plasma from sex- and age-matched healthy control mice was used to determine baseline protein expression. Source data for these analyses is provided in **Table S7**. We first visualized the distribution of all assessed markers with a principal component analysis which showed that aEGFRvIII CAR- and aEGFRvIII-SGRP CAR-treated plasma segregated the furthest from vehicle controls (**Fig. 4A**). To examine the peripheral immune response associated with the superior aEGFRvIII-SGRP CAR treatment, we performed differential expression analysis against aEGFRvIII CAR monotherapy (**Fig. 4B**), which showed significant enrichment of CCL3 and IL13 in aEGFRvIII-SGRP CAR-treated plasma (adj. *P* < 0.001 and adj. *P* < 0.05, respectively). Conversely, CD27 was significantly increased in aEGFRvIII CAR-treated plasma (adj. *P* < 0.05). To identify additional immune markers associated with aEGFRvIII CAR +/- SGRP, we looked for differences between SGRP-secreting CARs (aEGFRvIII-SGRP CAR vs aCD19-SGRP CAR) or non-SGRP-secreting CARs (aEGFRvIII CAR vs aCD19 CAR) across all 92 markers (**Fig. 4C**). This analysis revealed significant differences in the expression of CCL3, CXCL1, CXCL8, GZMA, TNFRSF21, TNFSF14 and VEGFA between SGRP-secreting CAR T cell treatments. Moreover, it showed significant differences in CD5, CXCL1, GZMA, IFNG, KLRD1, PGF and TNFSF14 expression between non-SGRP-secreting CAR T cell monotherapies. Collectively, these analyses identified CCL3 as the key plasma marker associated with aEGFRvIII-SGRP CAR treatment response out of all 92 markers studied. T cell-activation markers including GZMA and IFNG were found to be similarly highly expressed in aEGFRvIII CAR and aEGFRvIII-SGRP CAR plasma. Notably, CCL3, a myeloid chemoattractant, may have diverse roles in shaping peripheral responses to local anti-tumor therapy. A study investigating the role of CCL3 in driving cellular responses in tumor-draining lymph nodes in the priming phase of an anti-tumor response found that CCL3 recruits myeloid cells and drives dendritic cell (DC)-mediated T cell proliferation and differentiation in a CT26 colon tumor mouse model^49^. Although NSG mice carry defective DCs, a potential increase in CCL3-driven CAR T cell and DC/myeloid cell interactions could hint at a CD47-independent myeloid-modulating effect of aEGFRvIII-SGRP CAR T cell therapy.

**Figure 4.**
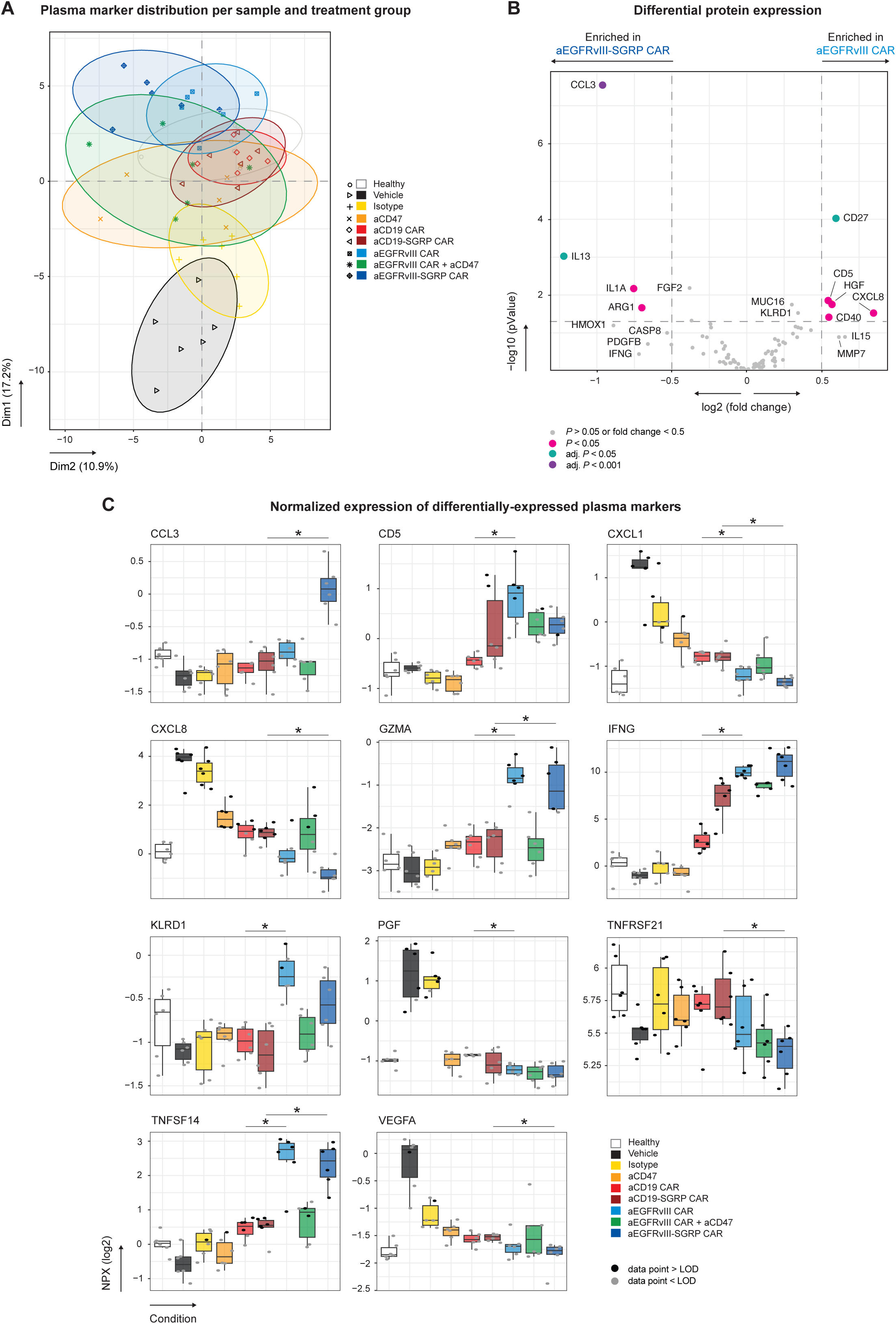
Local anti-EGFRvIII-SGRP CAR T cell therapy induces a potent inflammatory mediator response characterized by an increase in GAM chemoattractants. **A,** Non-metric multidimensional scaling (NMDS) plot of 92 examined soluble proteins in mouse plasma determined by proximity extension assay. Each mouse is represented by one data point, with symbols and colors differentiating healthy controls (healthy, white; n = 6), mock-injected (vehicle, black; n = 6), mouse IgG1-treated (isotype, yellow; n = 6), anti-human CD47-treated (aCD47, orange; n = 6), aCD19 CAR-treated (red; n = 6), aCD19-SGRP CAR-treated (dark red; n = 6), aEGFRvIII CAR-treated (blue; n = 6), aEGFRvIII CAR + aCD47-treated (green; n = 6) or aEGFRvIII-SGRP CAR-treated; (dark blue; n = 6). All samples were collected approximately 24 h after the last local treatment. The ellipses represent the 95% confidence interval of each condition. Source data are provided in **Table S7**. **B,** Differential expression analysis of aEGFRvIII-SGRP CAR vs aEGFRvIII CAR treatment groups, showing significant enrichment of immune markers CCL3, IL13, and reduction of CD27 in aEGFRvIII-SGRP CAR. Significant differences in protein expression are represented by differently-colored data points: adj. *P* < 0.001, purple; adj. *P* < 0.05, teal; *P* < 0.05, pink; *P* > 0.05 or fold change < 0.5, light grey. **C,** Scatter box plots showing the normalized protein expression (NPX) of significant innate immune surrogate markers in plasma: CCL3, CD5, CXCL1, CXCL8, GZMA, IFNG, KLRD1, PGF, TNFRSF21, TNFSF14 and VEGFA. Each data point represents one mouse. Data points above the assay’s limit of detection (> LOD) are illustrated in black and those below (< LOD) are illustrated in grey. The boxes’ central line represents the median, with the 75^th^ percentile at the upper bound, the 25^th^ percentile at the lower bound and the whiskers representing all samples lying within 1.5 times the interquartile range. Statistics were calculated using two-sided Mann–Whitney-U tests for the comparisons of interest (aEGFRvIII CAR vs aEGFRvIII CAR + aCD47, aEGFRvIII CAR vs aEGFRvIII-SGRP CAR, aEGFRvIII CAR vs aCD19 CAR, aCD19-SGRP CAR vs aEGFRvIII-SGRP CAR, aCD19 CAR vs aCD19-SGRP CAR, aEGFRvIII CAR + aCD47 vs aEGFRvIII-SGRP CAR, aCD47 vs aEGFRvIII CAR + aCD47) with Benjamini-Hochberg correction. Only significant statistics are shown.

### Anti-EGFRvIII-SGRP CAR T cells elicit an early rejection of EGFRvIII-mosaic GBM

To characterize the response of GBM xenografts to aEGFRvIII-SGRP CAR therapy, we collected spleen and brain tissue at an intermediate time point (7 days after the 2^nd^ treatment dose, 21 days post tumor implantation) for histological and immunofluorescence workup (**Fig. 5A**). Tumor size assessment calculated from H&E-stained brain histological sections confirmed the persistence of intracerebral tumor in all but aEGFRvIII-SGRP CAR-treated animals (**Fig. 5B**, and **Fig. 5C**). Only tumor remnants could be discovered in the brains of aEGFRvIII-SGRP CAR-treated animals at this time point, precluding us from a detailed multidimensional analysis of the immune TME in this condition despite whole-brain sectioning and staining. However, using anti-human CD3 IHC and applying an earlier time point (day 13 post tumor implantation) for the collection of aEGFRvIII-SGRP CAR-treated brains (**Fig. S6A**), we confirmed the persistence of CAR T cells in tumor core and tumor rim in all CAR treatment/controls (**Fig. S6B** and **Fig. S6C**).

**Figure 5.**
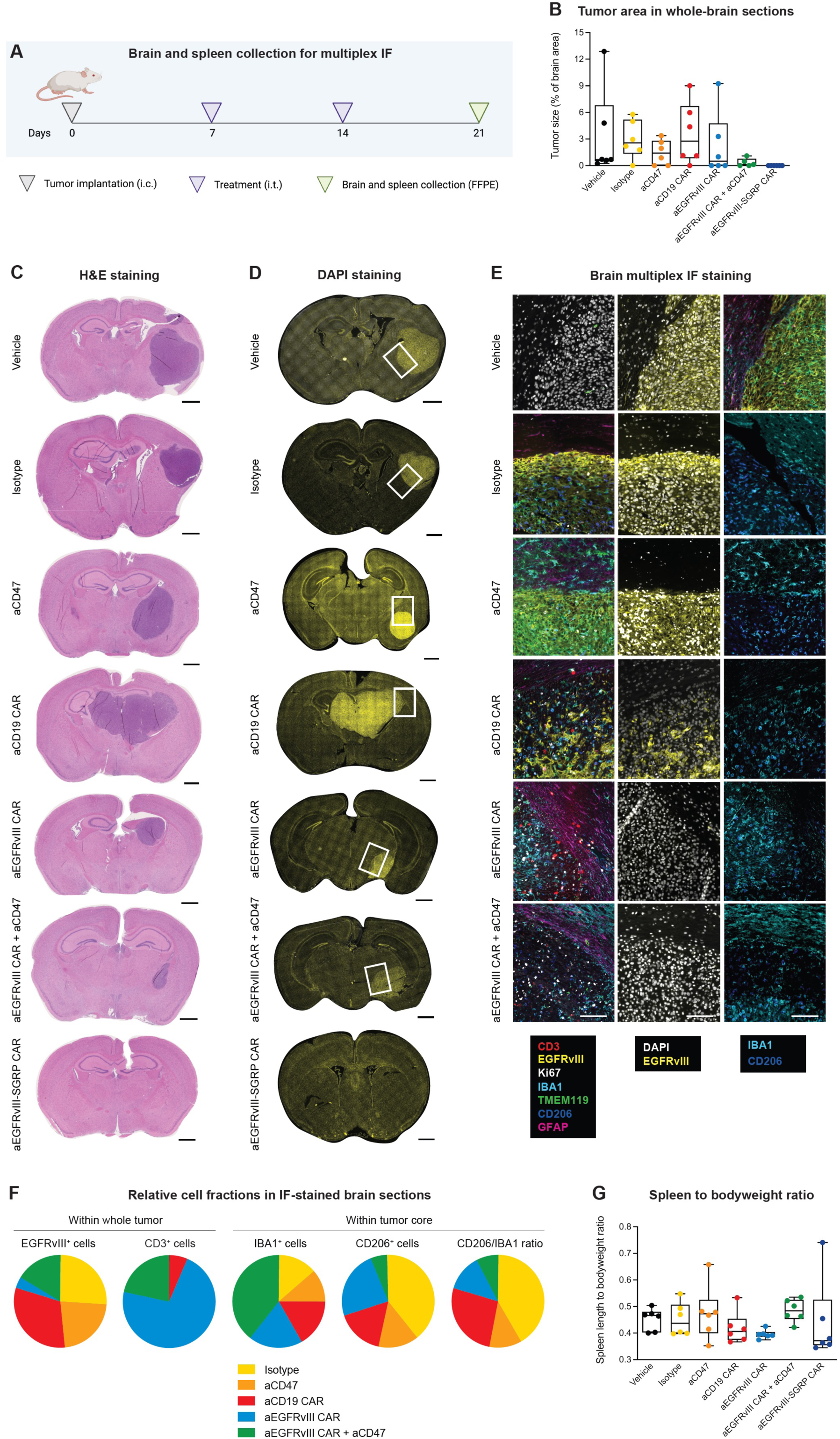
EGFRvIII-specific CAR T cells persist within the tumor and anti-EGFRvIII-SGRP CAR T cells eradicate EGFRvIII-mosaic GBM in histological brain sections. **A,** Brain and spleen collection at an intermediate post-therapeutic time point for multiplex immunofluorescence. **B,** Histomorphological brain tumor size assessment on day 21 post tumor implantation; n = 6 brains per condition, n = 5 brains in the aEGFRvIII CAR + aCD47 group. All comparisons were non-significant using one-way ANOVA with Dunnett’s multiple comparison tests. **C,** H&E-stained sections of representative tumor-burdened brains on day 21 post tumor implantation and 7 days after the second treatment dose. Frontal sections of cerebrum from different experimental groups from top to bottom: (1) mock-injected, vehicle, (2) mouse IgG1 (MOPC-21), isotype, (3) anti-human CD47 (B6.H12), aCD47, (4) aCD19 CAR, (5) aEGFRvIII CAR, (6) aEGFRvIII CAR + aCD47, and (7) aEGFRvIII-SGRP CAR; Scale bars: 1000 µm. **D,** DAPI-stained stitched assemblies of brain sections of experimental groups used for subsequent immunofluorescence multiplexing; Scale bars: 1000 µm. **E,** Representative micrographs of multiplexed immunofluorescence 4i assessment from different experimental conditions. *Left column:* overlays of 7 TME/tumor and proliferation markers: CD3 (red) for grafted CAR T cells, EGFRvIII (yellow) for target antigen-positive tumor cells, Ki67 (white) for cell proliferation, IBA1 (cyan) for mouse microglia and macrophages, TMEM119 (green) for mouse microglia, CD206 (dark blue) for protumoral GAMs, and GFAP (pink) for reactive astrocytes. *Center column*: EGFRvIII staining overlaid with DAPI, showing eradication of EGFRvIII^+^ cells by any EGFRvIII-specific CAR treatment paradigm. *Right column:* a qualitative assessment of CD206 (blue) protumoral GAMs overlaid with IBA1 (cyan) microglia and macrophages. Scale bars: 50 µm. **F,** Pie charts displaying relative comparisons of the percentage of marker-positive cells per all cells within the tumor or in the tumor core across experimental conditions. Two slides per condition were assessed creating a ratio between overall positive/negative cells. No tumors were detected in any of the aEGFRvIII-SGRP CAR-treated brains analyzed. Raw data are provided in **Table S8**. **G,** Assessment of spleen length to bodyweight ratios on day 21 post tumor implantation across all conditions.

Multiplexed immunofluorescence analysis focused on GAMs (IBA1, TMEM119, CD206), astrocytes (GFAP), CAR T cells (CD3) and tumor surface/proliferation markers (EGFRvIII, Ki67) within and surrounding brain tumor regions (**Fig. 5D** and **Fig. 5E**). The quantification of post-therapy EGFRvIII expression confirmed the selective elimination of EGFRvIII^+^ tumor cells after any aEGFRvIII-specific treatment, accompanied by persisting EGFRvIII^-^ tumor cells, except after aEGFRvIII-SGRP CAR treatment, where tumors were fully eradicated (**Fig. 5F**). CD3^+^ cells were found in all CAR treatment/control paradigms, albeit in higher relative numbers in aEGFRvIII CAR-treated brains (**Fig. 5F**). This substantiated the presence of tumor antigen as a driver of CAR T cell tumor-infiltration and persistence. Furthermore, the analysis of GAM markers revealed an overall increase in IBA1^+^ myeloid cell influx in aEGFRvIII CAR + aCD47-treated tumors (**Fig. 5F**) and a reduction of CD206^+^ cells in aCD47-treated tumors, consistent with a previous study^24^ (**Fig. 5F**).

Alternatively, we showed that anti-mouse CD68 IHC of aEGFRvIII-SGRP CAR-treated brains on day 13 post tumor implantation (compared to day 21 for all other conditions; **Fig. S7A**), had a more pronounced proinflammatory GAM activation at sites of tumor scarring (**Fig. S7B** and **Fig. S7C**). Taken together, these results suggest that constitutively secreted, paracrine-delivered SGRP into the TME is superior to antibody-based CD47 blockade for effective and sustained GAM modulation leading to a better anti-tumor response *in vivo*. Notably, despite its potent anti-tumor efficacy, aEGFRvIII-SGRP CAR showed no evidence of systemic cytokine toxicity as assessed by mouse survival (**Fig. 3C**), longitudinal monitoring of clinical signs and spleen size measurement (**Fig. 5G**).

### Anti-EGFRvIII-SGRP CAR treatment induces GAM-mediated tumor uptake and proinflammatory GAM conversion

To further elucidate the effect of the aEGFRvIII-SGRP CAR treatment in the iTME, we performed a focussed spectral FC analysis of myeloid cells shortly after i.t. treatment with CAR T cells (**Fig. 6A**). We designed and optimized a myeloid-targeted panel consisting of 17 surface markers, 1 intracellular marker and 2 fluorescent protein reporters (tagging grafted BFP2^+^ tumor cells and mCherry^+^ CAR T cells; **Table S5**). In comparison to vehicle control, both aEGFRvIII CAR and aEGFRvIII-SGRP CAR conditions triggered a rapid influx of CD45^+^ cells (**Fig. 6C**, upper row). We confirmed the presence of mCherry^+^ CAR T cells (**Fig. 6C**, middle row) and mTagBFP2^+^ tumor cells in the CD45^-^ gate and found that both CAR treatments reduced tumor cell burden at this stage (**Fig. 6C**, lower row).

**Figure 6.**
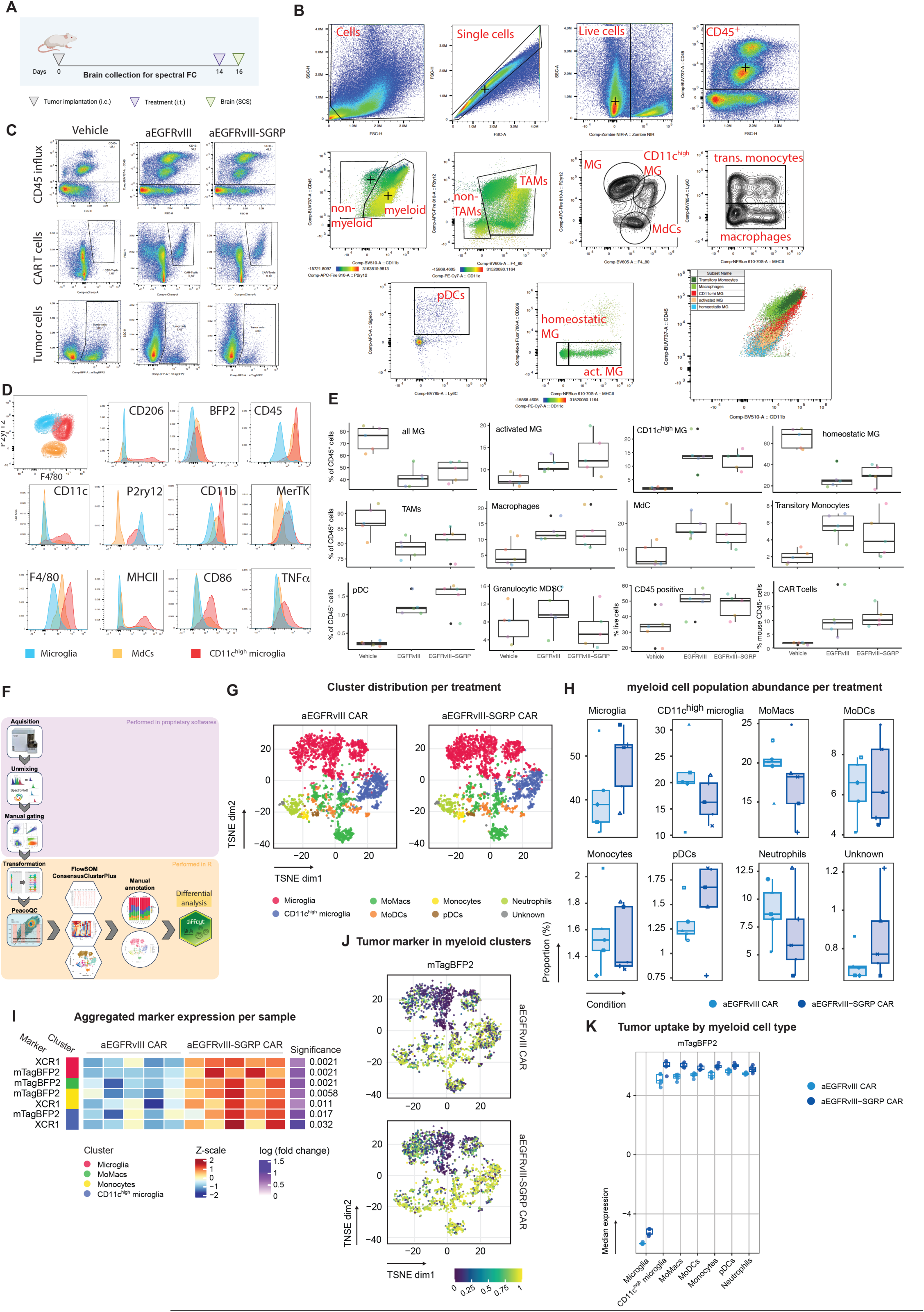
Anti-EGFRvIII-SGRP CAR treatment leads to increased myeloid cell-mediated tumor phagocytosis. **A,** Brain tumor collection at an intermediate post-therapeutic time point for multispectral flow cytometry. **B,** *Upper row:* Pseudocolor plots of a representative, conventional sample gating strategy to yield live (Zombie NIR^-^), CD45^+^ or CD45^-^ singlets. *Middle row, from left to right:* CD45^+^ cells were further subgated into CD11b^+^ myeloid and CD11b^-^ non-myeloid populations (the color bar insert represents MFI values of P2ry12-expression overlay on the plot); next, TAMs and non-TAMs were defined based on F4/80 expression (the color bar insert represents MFI values of CD11c-expression overlay on the plot); the density plot depicts different subsets within the TAM cluster: MG, CD11c^high^ MG, and MdCs; MdCs were further subdivided into transitory monocytes and macrophages based on Ly6C expression. *Lower row, from left to right*: within the non-TAM gate, pDCs were differentiated based on SiglecH expression; within the MG population, homeostatic MG and activated MG were differentiated based on MHCII expression; overlay backgating plot of the different identified myeloid populations within the CD45^+^ CD11b^+^ population. **C,** Representative pseudocolor plots derived from an individual single cell dissociation displaying differential amounts of CD45^+^ cells (upper row), mCherry^+^ CAR T cells (middle row) and mTagBFP2^+^ tumor cells (lower row) per therapeutic condition. **D,** Histograms of surface and intracellular marker expression within the TAM population, focussing on MG (blue), CD11c^high^ MG (red), and MdCs (orange). **E,** Scatter box plots of population frequency distribution across therapeutic conditions. Each dot represents an individual animal. Thick line represents the median value, box the interquartile range. Frequencies are represented as percentage within the CD45^+^ population, or, in case of CD45^+^ cells as percentage of live cells. CAR T cell frequency was measured within the mouse CD45^-^ gate. **F,** Flowchart visualizing the process of FlowSOM analysis and differential data analysis. **G,** t-distributed stochastic neighbor embedding (TSNE) plots depicting 8 main populations after initial merging and manual annotation from 15 populations originating from unbiased clustering (**Fig. S8**). **H,** Scatter box plots representing myeloid cell population abundance per therapeutic condition according to clusters identified in **G**. Each dot represents an individual animal (n=6); line at median, box represent interquartile ranges. **I,** Heatmap of aggregated marker expression per individual sample depicting the significant changes in expression of mTagBFP2 as a surrogate for tumor cell uptake and XCR1. Cluster ID/color: left X-axis, log fold change and significance level: right x-axis. **J,** Heatmap overlay of mTagBFP2 expression on TSNE from panel **G** per therapeutic condition**. K,** Scatter boxplot of median mTagBFP2 expression across identified myeloid cell populations. Each dot represents measurements from an individual brain (n = 6 per condition). Statistics: One-way ANOVA with Tukey’s multiple comparison test.

We then specifically elaborated on early time point differences between GAM subsets. The myeloid immune landscape in tumor-induced neuroinflammation harbors predominantly tissue-resident microglia (MG) and blood-borne monocyte-derived macrophages (MdCs) and dendritic cells (DCs)^50,51^. The advent of technologies that allow for high-dimensional analysis on a single cell level (scRNAseq, CyTOF, spectral FC) have added numerous flavors to MG and MdCs in health and disease ^52–55^, and established surface marker combinations that better define individual immune cell populations, therefore avoiding cross-contamination leading to ambiguous conclusions. To firmly differentiate MG from MdCs, we supplemented the classically used surface marker combination of CD45^int^CD11b^hi^ with the purinergic receptor P2ry12, as it has been shown to be highly specific for MG in health^56^ and in the tumor context^50^. Importantly, P2ry12 has been found to be downregulated during LPS-mediated MG activation^57^ and in neurodegeneration in the recently described disease-associated MG (DAM) population, which could potentially hamper our resolution of MG subpopulations ^53,54^. Using conventional analysis and gating-strategy (**Fig. 6B**), three main MG populations were resolved within TAMs: the MG gate (CD45^int^CD11b^hi^P2ry12^hi^) could by divided into ‘homeostatic’ MG characterized as CD45^int^CD11b^hi^P2ry12^hi^F4/80^int^MHCII^-^ and ‘activated’ MG which additionally express MHCII (**Fig. 6B**). Furthermore, we identified a P2ry12^int^CD11c^hi^ MG population, and a P2ry12^-^F4/80^+^ MdC-derived population that was subdivided into Ly6C^+^ transitory monocytes and Ly6C^-^ terminally differentiated macrophages (encompassing varying MHCII expression), according to^58^.

CD11c^hi^ MG arose only upon CAR T cell treatment and were undetectable in non-CAR-treated mice (**Fig. 6E**). Multiple studies described MG subpopulations expressing CD11c (*Itgax*) in healthy^52^ and diseased mouse brain^59–61^. By Seurat-based integration of six CNS-macrophage focused scRNAseq datasets, Silvin et al.^54^ recently incorporated the transcriptome of healthy (fetal, adult and old) and diseased MG/macrophages into one universe, termed macrophage-verse. Projection of the transcriptomic signature of CD11c^+^ MG sampled from the corpus callosum (formerly termed “fountain of MG”) of postnatal mice^59^ (p7 CD11c^+^ MG) showed strong overlap with the projected and further deconvoluted disease-associated microglia (DAM) described by Keren-Shaul ^53^. Furthermore, p7 CD11c^+^ MG share a remarkably coherent signature with p4/p5-age-restricted axon-tract-MG (ATM)^52^ and were shown to be a specialized phagocytic MG subset localized within the corpus callosum essential to physiological myelogenesis. It seems that CD11c^+^ MG mark a conserved MG subset that are spatially and temporarily restricted to an early postnatal phagocytic MG phenotype within white matter in health and reappear during neuroinflammation with a neuroprotective role^53^.

Thus, DAMs display higher expression of CD11c, CD45, F4/80, MHCII, CD206 and lower expression of Cx3cr1 and P2ry12 compared to ‘homeostatic microglia’. The identified CD11c^hi^ MG displays this exact surface DAM signature (**Fig. 6D**) and - in line with p7 CD11c^+^ MG, ATM and DAMs - functionally turns out to be a phagocytic (shown by mTagBFP2-positivity) MG population (**Fig. 6D**). CD11c^hi^ MG are also highly positive for CD206 (**Fig. 6D**), a marker that has classically been used to define the M2-polarized state of macrophages, but recently was shown to correlate strongly with phagocytosing macrophages^62^. We therefore suggest that the conserved and phagocytic program subsumed as DAM-signature likewise reappears in the tumor context. Moreover, this seems to be in a T-cell dependent manner as CD11c^hi^ MG are restricted to CAR-T-treated conditions in a NSG background lacking any lymphocytes (**Fig. 6E**).

Moreover, we observed a CAR-T-cell-dependent significant increase of transitioning monocytes and terminally differentiated macrophages (**Fig. 6E**), supporting the recent report on the dependency on T-cell-mediated IFNγ for the monocyte-to-phagocyte transition^58^.

Hence we would like to note that the gating strategy based on CD45^int^CD11b^hi^ expression should be supplemented by further MG-specific markers such as P2RY12 to increase resolution of MG-subsets and avoid cross-contamination of macrophage population in neuroinflammation (**Fig. 6C**). The above mentioned T-cell restricted immune cell appearance was also observed for the CD45^+^CD11b^-^Ly6c^+^SiglecH^+^ pDC population (**Fig. 6C, E**).

We alternatively performed a multiparametric analysis combined with FlowSOM meta-clustering of GAMs (**Fig. 6F**). Initial clustering identified 15 meta-clusters (TSNEs/UMAP in **Fig. S9-p1-A**) that were characterized by differential lineage marker expression (**Fig. S8-p1-B,C**). Cluster frequencies between aEGFRvIII CAR and aEGFRvIII-SGRP CAR treatment conditions were not different (**Fig. S8-p1-D**). Subsequently, these clusters were merged into 8 myeloid subsets including MG, activated MG, monocyte-derived macrophages (MoMacs), monocyte-derived dendritic cells (MoDCs), monocytes, plasmacytoid dendritic cells (pDCs), neutrophils, and unknowns (**Fig. 6G**, TSNE plots and UMAP in **Fig. S8-p2-A,B**), which exhibited similar population frequencies among CAR-T-treated conditions (**Fig. 6H**). We next focused on the overall most abundant phagocyte population, MG, which expressed P2ry12, F4/80 and CD11c in varying levels as seen in the conventional analysis (**Fig. 6C**). Further MG-specific sub clustering revealed 3 major populations (homeostatic MG, MHCII^high^ activated MG, and CD11c^hi^ MG, TSNE in **Fig. S8-p3-A**), characterized by marker profiles outlined in **Fig. S8 B-E**). While population frequencies between aEGFRvIII and aEGFRvIII-SGRP CAR-treated animals were not different (**Fig. S8-p3-F**), differential abundance analysis among conditions showed significantly increased mTagBFP2 expression in most myeloid cells in aEGFRvIII-SGRP CAR-treated brain tumors (expression heatmap in **Fig. 6I-K, S8-p3-G,** individual heatmaps of all assessed markers in **Fig. S8-p4**). Furthermore, XCR1, a specific classical dendritic cell marker involved in antigen presentation^63^, was increased in MG and monocytes upon aEGFRvIII-SGRP CAR treatment. Taken together, this early time point analysis after aEGFR-SGRP CAR intratumoral treatment hints towards a pronounced induction of tumor cell phagocytosis, and showcases the strong myeloid reaction induced by intratumoral CAR T cell application.

### Anti-EGFRvIII CAR T cells outperform anti-CD19 CAR T cells in co-cultures with patient-derived single-cell suspensions

To demonstrate the efficacy of aEGFRvIII-SGRP in a translational setting, we co-cultured CAR T cells with EGFRvIII^+^ GBM patient-derived single-cell suspensions and performed a multidimensional image-based analysis in 5 patients (**Fig. S9, S10A**). The clinical characteristics of included patients are listed in **Table S9**. CAR T cells and patient-derived GBM cells were not matched in this setup, raising the potential of alloimmunological phenomena. EGFRvIII status was determined by a custom qPCR assay (**Fig. S9**). Readout included an assessment of NESTIN^+^ tumor cells and EGFRvIII^+^ tumor cells, as well as CD14^+^ immune cell counts (**Fig. S10B**). In this setting, a direct benefit of aEGFRvIII-SGRP CAR over aEGFRvIII CAR was not appreciated. As expected, all EGFRvIII-specific CARs significantly reduced the number of EGFRvIII^+^ cells compared to CD19-specific control CARs (**Fig. S10C**). Moreover, aCD19-SGRP CARs induced moderate antigen-independent killing of both EGFRvIII^+^ and Nestin^+^ tumor cells, presumably due to a potential SGRP-mediated effect or constitutive secretion of IFNγ (**Fig. S10D)**. Further, CD14^+^ myeloid cells were reduced in the aEGFR-CAR treated conditions, presumably due to a short term bystander effect by CAR-mediated cytokine release (**Fig. S10E**). The fact that no differential effect between aEGFRvIII and aEGFRvIII-SGRP was detected in this experiment could be explained by the use of frozen single-cell suspensions with reduced overall viability and subsequent functional deficits in myeloid cell composition.

### Anti-CD19-SGRP CAR T cells have superior efficacy over conventional aCD19 CAR T cells in a peripheral lymphoma xenograft model

To demonstrate a potential benefit of immune TME-targeted secretion of SGRP in another solid tumor context, we further assessed the efficacy of aCD19-SGRP CAR T cells against CD19^+^ lymphoma xenografts. In contrast to locoregional injection as the preferred application route in brain tumors, we considered a systemic approach mirroring CAR T cell treatments currently performed against leukemia^64^. CD19^+^ Raji cells were injected s.c. in the right flank of NSG mice followed by a single dose of systemic CAR T cell infusion 3 days post tumor implantation. Animals were monitored for survival analysis and volumetric tumor burden quantification (**Fig. 7A**). The therapeutic setup consisted of target-specific CAR T cells (aCD19 CAR or aCD19-SGRP CAR) or -unspecific CAR T cells (aEGFRvIII CAR or aEFGRvIII-SGRP CAR) serving as controls (**Fig. 7B**). All aCD19 CAR-treated animals had a significant survival benefit compared to either vehicle or target-unspecific CAR controls (**Fig. 7C**). Strikingly, aCD19-SGRP CAR treatment resulted in the longest survival benefit (20% overall survival) with one cured animal (undetectable tumor), trending towards a superior response compared to conventional aCD19 CAR application (*P* = 0.0731; **Fig. 7C** and **Fig. 7D**). Thus, the contribution of SGRP-mediated innate immune modulation is relevant in solid cancers other than GBM. This data demonstrates that our approach could potentially be used for the treatment of other types of cancer where myeloid immune suppression but also tumor heterogeneity are major hurdles to effective CAR T cell therapy.

**Figure 7.**
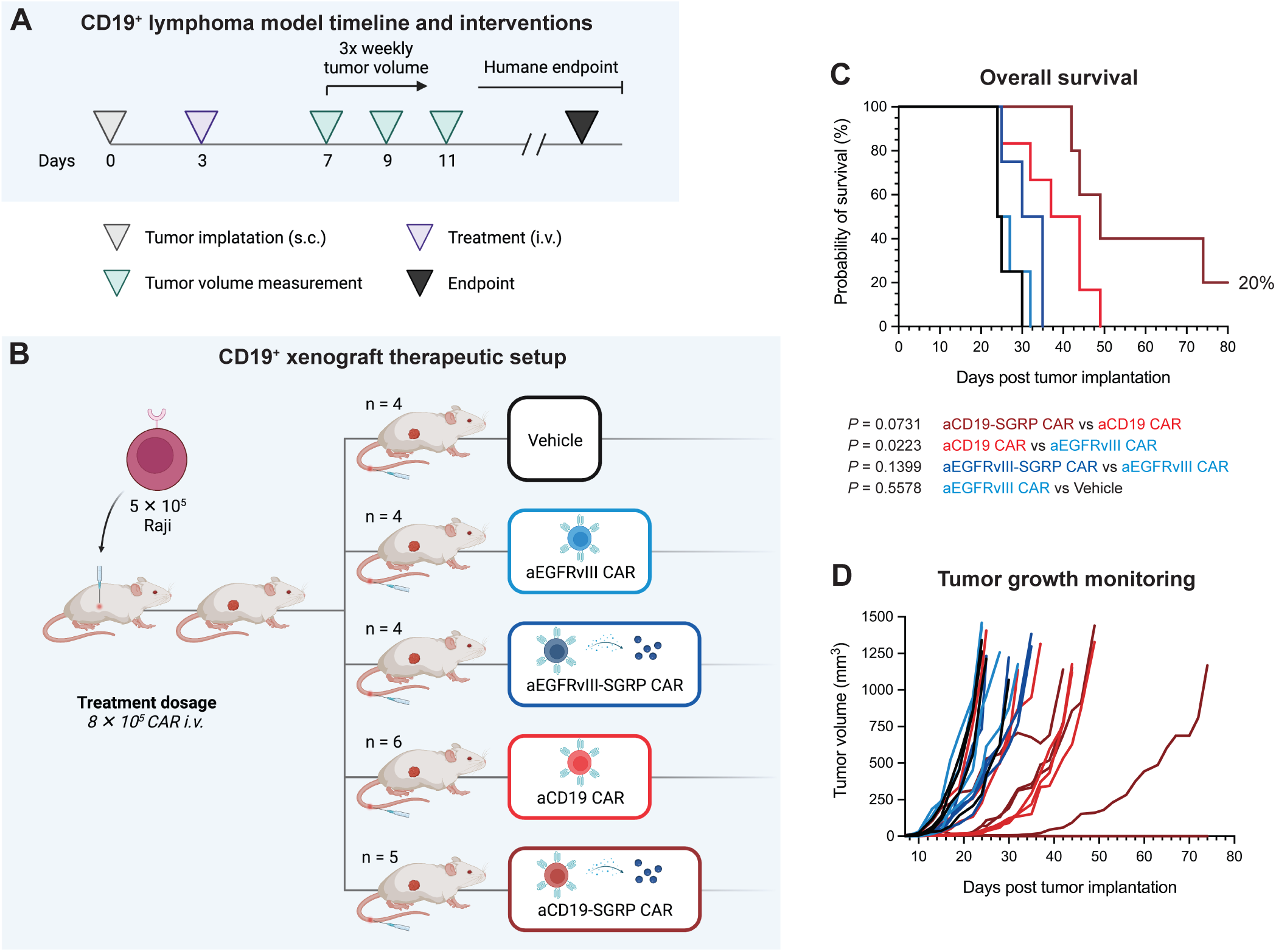
Systemic anti-CD19-SGRP CAR T cell therapy delays tumor growth and improves survival in a CD19^+^ lymphoma xenograft model. **A,** Experimental setup and timeline of interventions of peripheral CD19^+^ lymphoma model treated with systemic CAR T cell infusions. Three days after tumor implantation in the right flank, mice were treated i.v. with CAR T cells, followed by 3 times weekly tumor volume assessment and clinical scoring. Mice were sacrificed upon reaching the humane endpoint. **B,** Overview of experimental groups/therapeutic conditions and treatment dosages. **C,** Kaplan–Meier plot of overall survival (in days). Log-rank tests were used to compare the indicated treatment/control groups. **D,** Tumor volume measurements in mm^3^ of individual animals over time (in days post tumor implantation).

## Discussion

Myeloid suppression and tumor heterogeneity are the main factors for CAR T cell therapy failure against GBM. In this study, we show the superior efficacy of armored aEGFRvIII-SGRP CAR T cells in EGFRvIII-mosaic GBM orthotopic xenograft and patient-derived explant models, offering a potential approach to treat EGFRvIII^+^ GBM. This novel CAR will now be tested in a first-in-human clinical trial (NCT ID pending) for patients with recurrent glioblastoma by neoadjuvant locoregional application of CAR T cells. Moreover, armoring aCD19 CARs with SGRP significantly improved efficacy compared to conventional aCD19 CARs in a subcutaneous lymphoma model, highlighting the strong effect of paracrine GAM modulation by high-affinity CD47-SIRPɑ disruption in other tumor entities and application routes. If our results are clinically validated in recurrent GBM, this approach could provide a unique option for the treatment of primary EGFRvIII^+^ GBM tumors upfront, eventually reducing the reliance on cytotoxic chemotherapy and irradiation and significantly improving patient outcomes. It is important to note that EGFRvIII is one of the few tumor-specific antigens in GBM known to date and one that has been demonstrated to be safe as a CAR T cell target in clinical GBM studies^10,34^. Yet, antigen escape might pose a threat to all EGFRvIII-targeted treatments^65^, underscoring the need for careful patient stratification using reliable pre-therapeutic plasma^66^ or tissue biopsy analysis to confirm the presence of this oncogenic mutation also in recurrent GBM. Importantly, recent studies using more sensitive methods of detection, suggest that even up to 70% of GBM patients express EGFRvIII^66^.

Nevertheless, paracrine secretion of SGRP, which has a high affinity to CD47 on tumor cells as well as normal cells, might pose a risk for toxicity. While this was clinically absent in the mouse models applied in our study, potential exacerbation of SGRP-mediated hematologic toxicity such as induction of anemia^27^ needs to be carefully considered in first-in-human phase I studies. Locoregional application of CAR T cells for brain tumor treatment is superior to systemic infusion in terms of efficacy, prevention of preconditioning lymphodepletion and reduced systemic side effects^67^. Although this was not assessed for ‘armored’ CARs in our study, the peripheral effect of leaking SGRP may be significant. While we did not measure SGRP levels in the plasma of treated animals, we found significant SGRP-related levels of the macrophage chemotactic cytokine CCL3 as a surrogate for a potential peripheral innate immune induction. This might be an indicator of a beneficial antitumoral response, but could also be related to potential adverse events that need to be carefully monitored in the clinical setting. Hence, studies assessing other local combinatorial CAR/immunomodulatory strategies such as a combination of EGFRvIII-CARs with intratumoral IL-12^68^ or with concomitant bispecific T-cell engager (BiTE) expression^69^ did not report increased systemic toxicity *in vivo*.

Direct paracrine secretion of high-affinity CD47 blockade might preferentially target the solid tumor cells in trans and impact autocrine T cell CD47 blockade to a lesser extent, shifting the balance towards GAM reprogramming. Further, the engineered CAR with strong co-stimulatory signals could equally contribute towards T cell viability/activity, and escape the potential effects of CD47 blockade by both antibodies or SGRP. Even though a small contribution of secreted SGRP could elicit minor therapeutic benefits as seen in the case of target-unspecific aCD19-SGRP CAR T cells against human GBM or when aEGFRvIII-SGRP against CD19^+^ tumor cells were used, the proximity of the effector CAR T cell to the iTME seems to be extremely important and speaks in favor of local CAR T cell delivery. Second, we treated the animals with anti-CD47 antibody clone B6H12 which might have lower efficacy than the antibody H5F9 used in clinical trials - leading to inferior antitumor activity. However, systemic anti-CD47 mono-treatment failed to yield substantial survival benefits in aggressive brain tumor models in several studies^70,71^. Third, we did not perform continuous local maintenance dosing of anti-CD47 antibodies which might mask optimal efficacy of anti-CD47 treatment. On the other hand, this might have aggravated the effect on CD47^+^ grafted CAR T cell persistence further. Recent studies of antagonizing CD47 by Sirpa-FC secreted by CAR T cells in peripheral lymphoma context support the rationale and efficacy of SGRP-mediated CAR T cell efficacy observed here in GBM ^72^.

While the *in vivo* results in several experimental series were convincing, our study has limitations that open room for further research and need to be considered in a potential clinical setting. While the direct paracrine secretion of SGRP into the TME was accompanied by a strong antitumoral, curative response in heterogeneous GBM, the combination of locoregional/systemic maintenance of anti-CD47 treatment with aEGFRvIII CAR T cells did not meet the expected synergistic efficacy. Several factors might explain why this combination failed. First, human T cells express substantial levels of CD47. Therefore, blockade of CD47 using high doses of anti-CD47 antibodies might mark the grafted T cells for host-mediated phagocytosis. Recently, a study by Becket et al. indeed found T cell fratricide when engineered CAR T cells expressing a high-affinity Sirpα variant as a chimeric receptor^73^. Moreover, CD47-deficient T cells were recently reported to be removed by dendritic cells via necroptosis^74^. In the multidimensional immunofluorescence analysis 7 days after the last treatment, we observed intratumoral CAR T cell reduction upon combining with anti-CD47 antibodies compared to aEGFRvIII CAR treatment alone, corroborating these observations. However, conventional IHC quantification did not reveal major differences between intratumoral CAR T cell conditions.

The expansion protocol in this study preferentially yielded CD4^+^ CAR T cells, and all *in vivo* experiments were performed using this subset with the described efficacy. The indirect killing effect of IFNγ has been described to be preferentially attributed to CD4^+^ CAR T cells, enabling target-negative tumor cell killing. Indeed, we found high levels of IFNγ secretion, but only SGRP-armored CARs were able to cope with a high tumor load of target-negative cells. Constitutively secreted IFNγ could cause the non-targeted tumor cell loss observed in our pharmacoscopy studies and its blockade needs to be considered when responsible for systemic toxicity. On the other hand, tumor cell bystander killing might be of high importance in the eradication of heterogeneous GBM cells.

While we did not directly demonstrate a pro-phagocytic effect of SGRP-induced macrophage-mediated tumor cell phagocytosis *in vitro*, we identified SGRP-related, increased uptake of tumor cells by CD11c^hi^ microglia and other myeloid subsets *in vivo*. Furthermore, SGRP induced expression of the chemokine XCR1 on activated microglia, previously attributed to the dendritic cell antigen presentation machinery^75^. Even if the impact of XCR1 upregulation on microglia in a NSG background on endogenous T cells cannot be validated, a pronounced induction of antigen presentation would be highly desirable, especially in the frame of a clinical trial. Direct local application of SGRP by e.g. mini osmotic pumps would provide additional insights into its effect on the immune TME. Taking into account that aCD19-SGRP CAR T cells in the brain tumor xenografts elicited only a moderate effect, SGRP by itself might in line with anti-CD47 act merely as a sensitizer of enhanced tumor cell phagocytosis and additional ‘eat-me’ signal would be needed to yield a durable antitumoral response. Further, a recent study of antagonizing CD47 by a high-affinity SIRPα secreted by CAR T cells in peripheral lymphoma context supports the rationale and efficacy of SGRP-mediated CAR T cell efficacy observed here in GBM^72^.

## Conclusion

In conclusion, we showed significant synergy of aEGFRvIII-SGRP CARs against orthotopic and patient-derived GBM models, credentialing this approach for a first-in-human clinical trial. Additionally, this work highlights the importance of paracrine, high-affinity CD47 blockade to relieve macrophage-mediated immunosuppression, which might not be achievable by systemic or episodic local application of anti-CD47 antibodies in the context of aggressive solid tumors. Potentiating other CAR T constructs with secretable SGRP might form an important platform in the armamentarium against solid tumors.

In summary, this work represents one of the first examples of GBM targeting by adoptive T cells with concurrent local paracrine GAM modulation in a single therapeutic modality.

## Supporting information

Supplementary protocol 1

Supplementary protocol 2

## Supplementary Figure Legends

**Figure S1.**
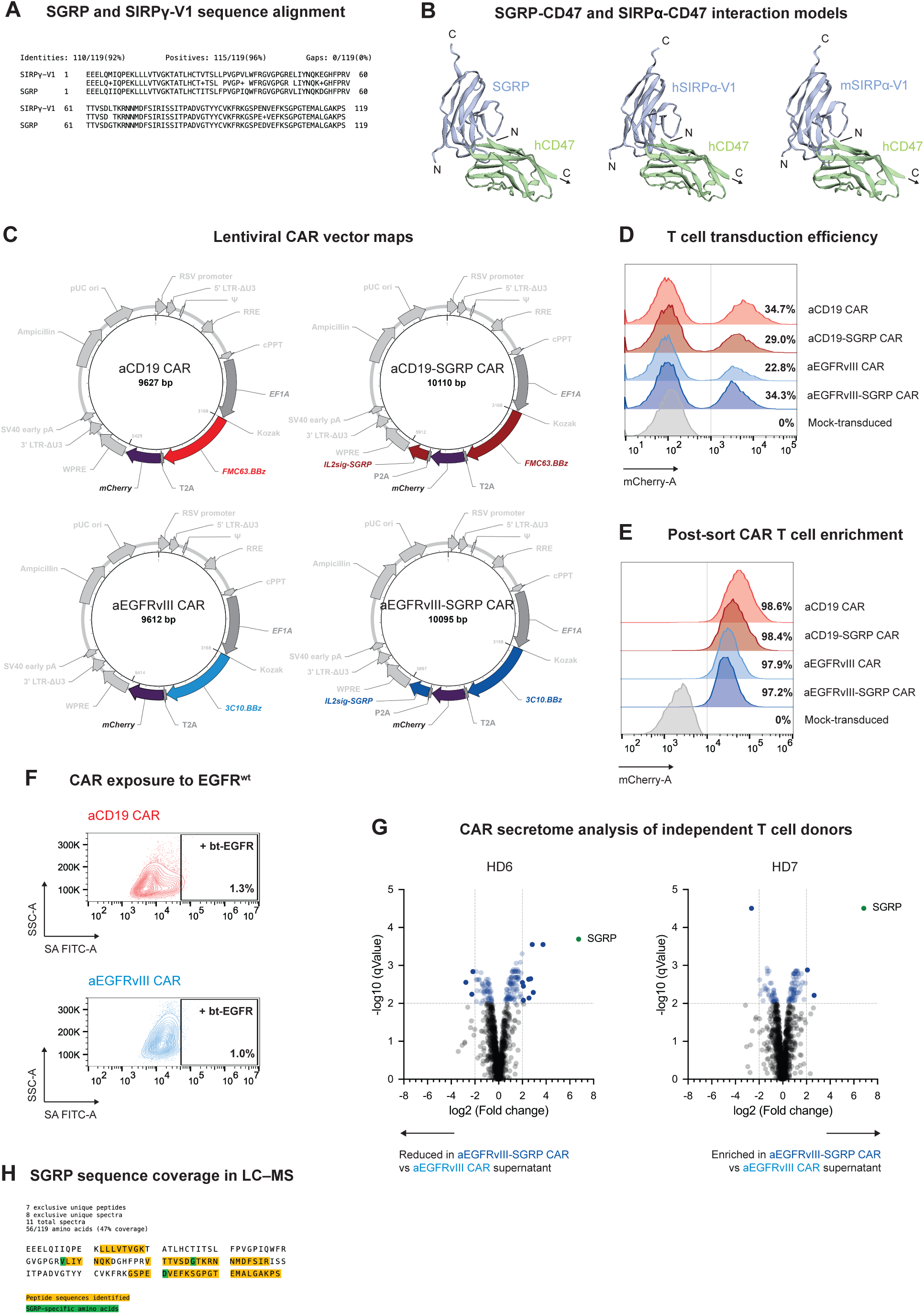
Overview of used CAR constructs, sequence alignments and LC–MS supernatant analysis. **A,** SGRP and homologous SIRPγ sequence alignment encompassing the whole length of SGRP (119 AA). **B,** AlphaFold^31,32^-generated *in silico* modeling displaying predicted protein-protein interactions of SGRP, hSIRPα-V1 and mSIRPα-V1 with hCD47. Amino acid sequence information regarding hCD47, IL2sig, hSIRPα-V1, hSIRPγ-V1, IL2sig-SGRP, mSIRPα-V1 and SGRP are listed in **Table S1**. **C,** Overview of lentiviral vector maps used in the study. Nucleotide sequences for aCD19 (FMC63.BBz), aEGFRvIII (3C10.BBz), aCD19-SGRP and aEGFRvIII-SGRP constructs are listed in **Table S2**. **D,** Representative T cell lentiviral transduction efficiency for the different CAR lentiviral vectors used, based on the percentage of mCherry^+^ T cells assessed by FC 4 days after transduction (gated on live single cells). Mock-transduced T cells served as controls. **E,** Post-sort enrichment of CAR T cells after sorting for mCherry^+^ cells. These were subsequently expanded and used for downstream experiments. **F,** aCD19 or aEGFRvIII CAR T cells were exposed to wt-EGFR using biotinylated (bt) EGFR recombinant protein. No binding of CARs to bt-EGFR by FC analysis of streptavidin (SA)-FITC could be detected. **G,** Volcano plots of healthy-donor (HD) specific differential supernatant secretome analysis by LC–MS of aEGFRvIII-SGRP CAR vs aEGFRvIII CAR T cells. Log2 (Fold change) indicates the mean expression level for each protein. Each dot represents one protein. The -log10 (qValue) represents the adjusted level of significance for each protein. SGRP (green dot) is highly enriched in aEGFRvIII-SGRP CAR T cell-conditioned media from both HDs assessed. The raw expression data is available in **Table S3**. **H,** Representation of SGRP sequence coverage in LC–MS. SGRP was identified by 7 exclusive unique peptides and 8 exclusive unique spectra covering 47% of the 119 AA sequence.

**Figure S2.**
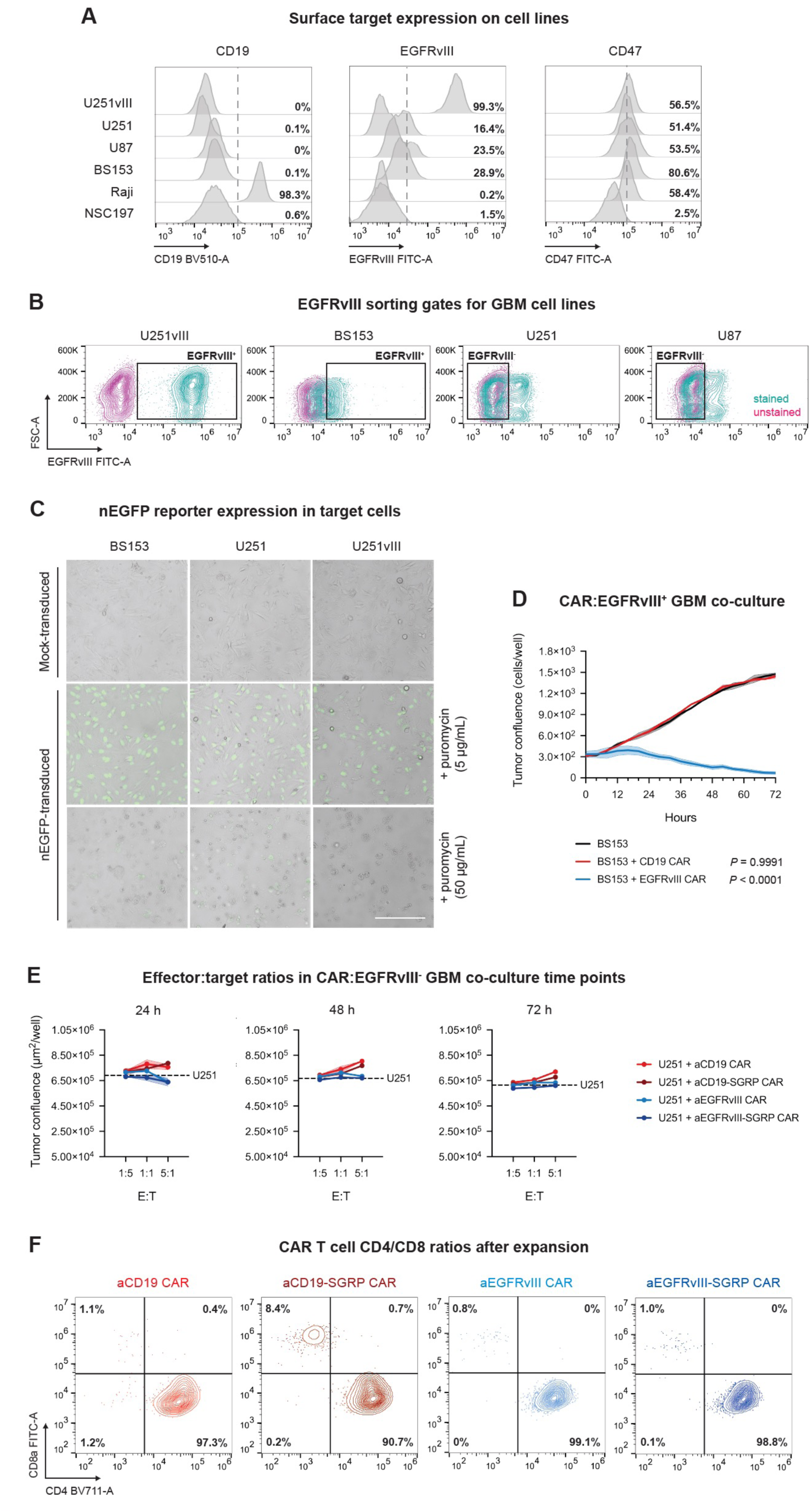
*In vitro* characterization of tumor cell lines, CAR T cell efficacy and CAR T cell CD4/CD8 phenotyping. **A,** FC representation of surface target expression of tumor cell lines U251vIII, U251, U87, BS153, Raji and control neural stem cell line NSC197. **B,** FC gates showing the EGFRvIII-stained (teal) populations sorted for subsequent *in vitro* and *in vivo* experiments encompassing EGFRvIII^+^ U251vIII or BS153 and EGFRvIII^-^ U251 or U87 cells. **C,** Cell culture microphotographs overlaying brightfield and fluorescent detection of nuclear (n)EGFP-reporter expression after lentiviral, puromycin-selective transduction of target cells BS153, U251 and U251vIII for later time-lapse co-culture experiments; Scale bar: 250 µm. **D,** 72 h time-lapse co-culture experiment of endogenously EGFRvIII^+^ BS153 cells with aCD19 or aEGFRvIII CARs, respectively. Tumor confluence (cells/well) is plotted over time. Curves represent the mean of duplicate measurements. Curve-adjacent shaded areas represent standard deviation. Differences between each co-culture compared to the ‘tumor alone’ control condition were analyzed using one-way ANOVA and Dunnett’s multiple comparisons tests. **E,** Tumor confluence (green nuclei per well) in co-cultures of EGFRvIII^-^ U251 with either aCD19, aCD19-SGRP, aEGFRvIII or aEGFRvIII-SGRP CAR T cells in 3 different effector-target ratios at 24, 48 and 72 h timepoints. Dashed line: U251 cell line alone. **F,** FC assessment of CD4/CD8 T cell phenotypes of aCD19, aCD19-SGRP, aEGFRvIII and aEGFRvIII-SGRP CAR T cells used in this study (gated on live singlets) at the time of experimental use. One representative healthy donor CAR T cell batch is shown.

**Figure S3.**
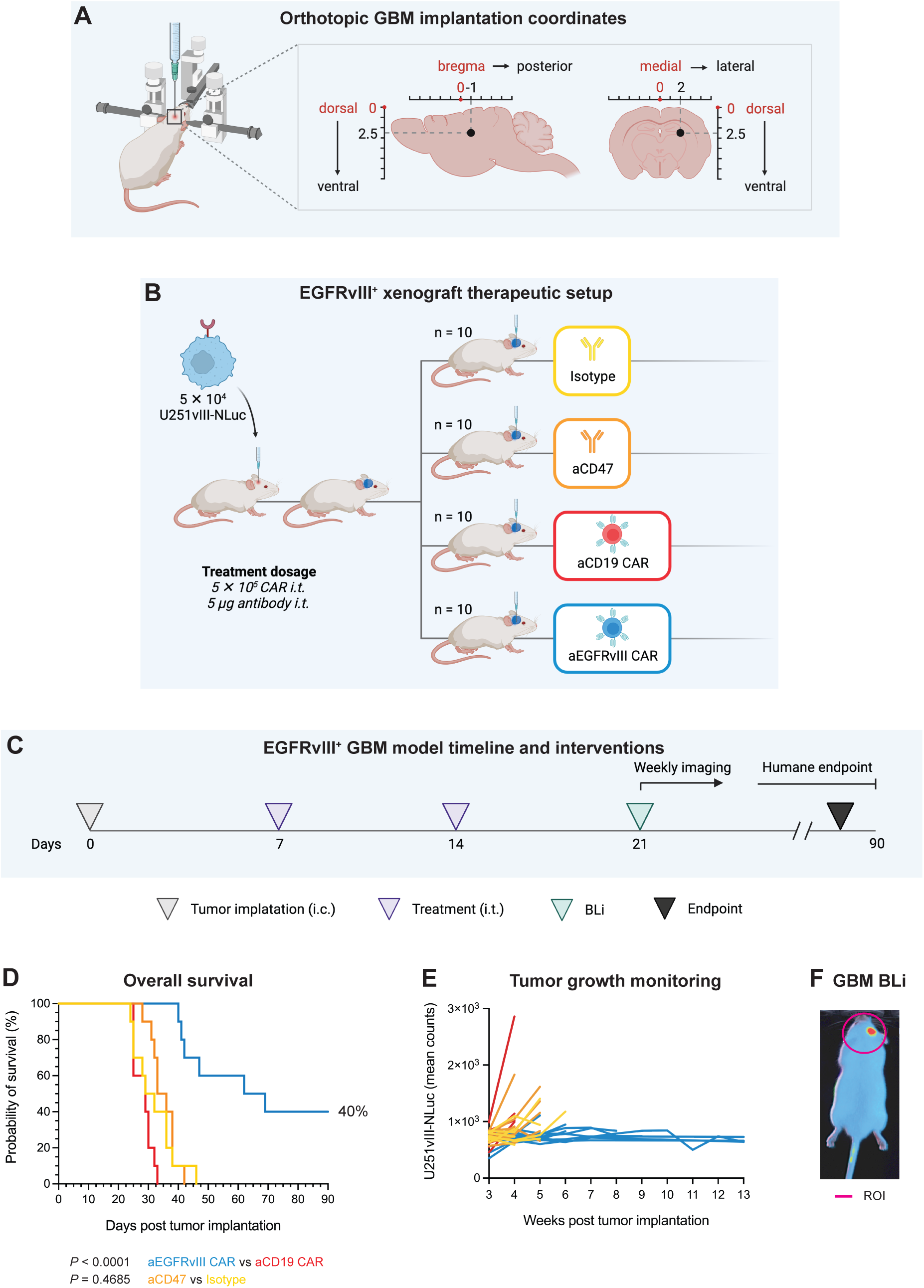
Setup and results of orthotopic EGFRvIII-homogeneous GBM *in vivo* experiments. **A,** Coordinates (in mm) of orthotopic GBM implantation in adult NSG mice. **B,** Overview of the experimental setup of the EGFRvIII^+^ xenograft GBM tumor model and subsequent monotherapeutic CAR or antibody treatments. **C,** Animals were treated with either i.t. CAR T cells or antibodies 7 and 14 days after orthotopic tumor implantation with U251vIII-NLuc tumor cells, followed by BLi and scoring until the humane endpoint was reached. **D,** Kaplan–Meier plot of overall survival (in days). Log-rank tests were used to compare the indicated treatment/control groups. **E,** Tumor progression in EGFRvIII^+^ xenografts was monitored using BLi time course imaging with FFz substrate (in weeks). **F,** Head ROI shape defined for all BLi measurements throughout the study.

**Figure S4.**
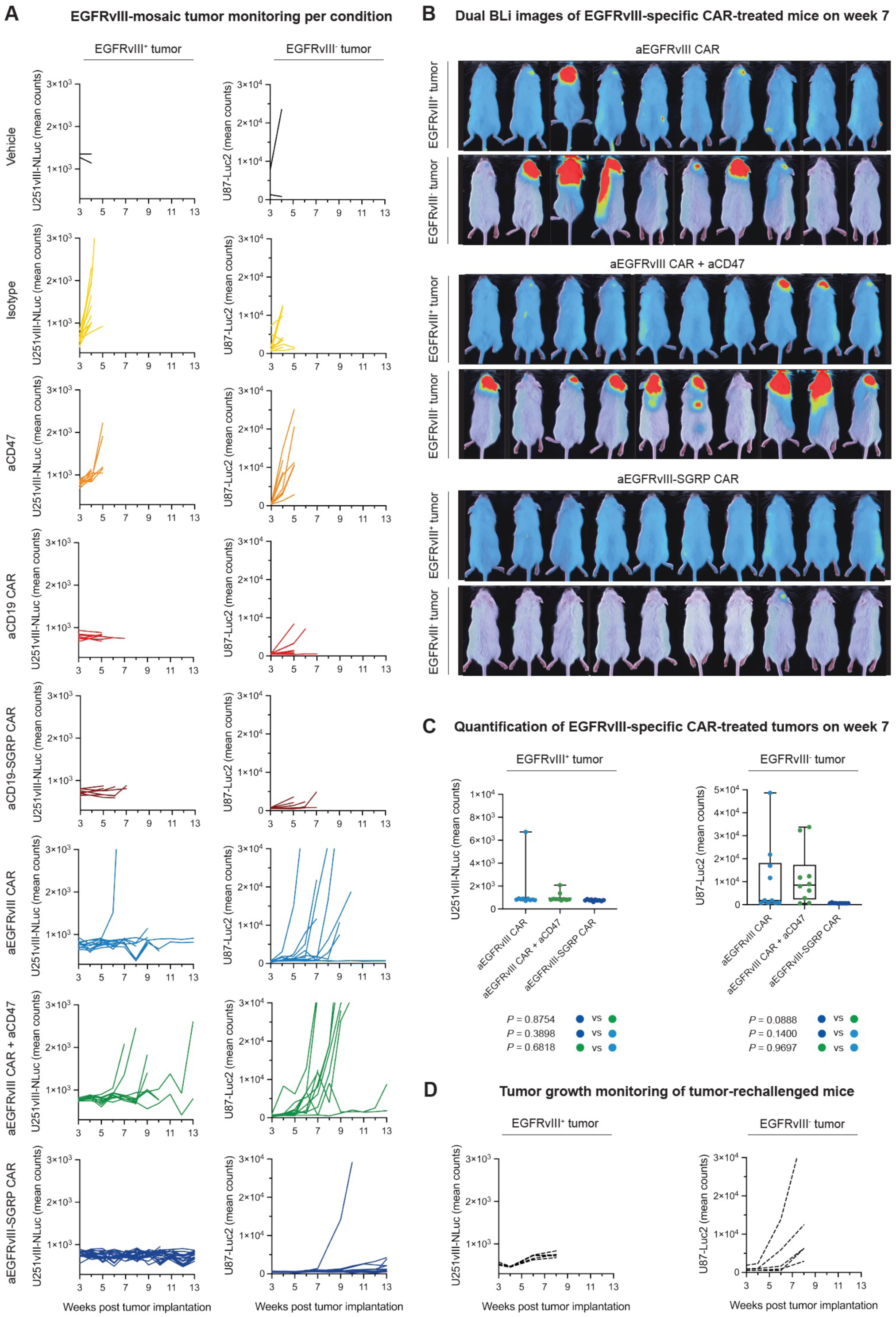
Overview of *in vivo* bioluminescence readouts. **A,** Cumulative differential monitoring in weeks by BLi for both grafted mosaic EGFRvIII^+^ and EGFRvIII^-^ tumors per experimental condition as outlined in Fig. 3A and Fig. 3B. U251vIII^+^ tumor growth was measured by luminescence elicited by FFz, whereas growth of U87vIII^-^ tumors was detected by D-luciferin luminescence. Curves end whenever the humane endpoint was reached. **B,** Representative overlay images of dual BLi studies at week 7 of aEGFRvIII CAR, aEGFRvIII CAR + aCD47 and aEGFRvIII-SGRP CAR treated animals highlighting the suppression of EGFRvIII^-^ tumors in aEGFRvIII-SGRP CAR treated animals. **C,** Quantification of BLi signal intensities (mean photon counts) as a surrogate for tumor burden for both EGFRvIII^+^ and EGFRvIII^-^ tumors from (**B**) at 7 weeks after tumor implantation. Each dot represents an individual mouse. Statistical comparisons were performed with one-way ANOVA with multiple comparisons corrections. **D,** Dual BLi monitoring of initially cured, mosaic tumor rechallenged mice in weeks after tumor rechallenge.

**Figure S5.**
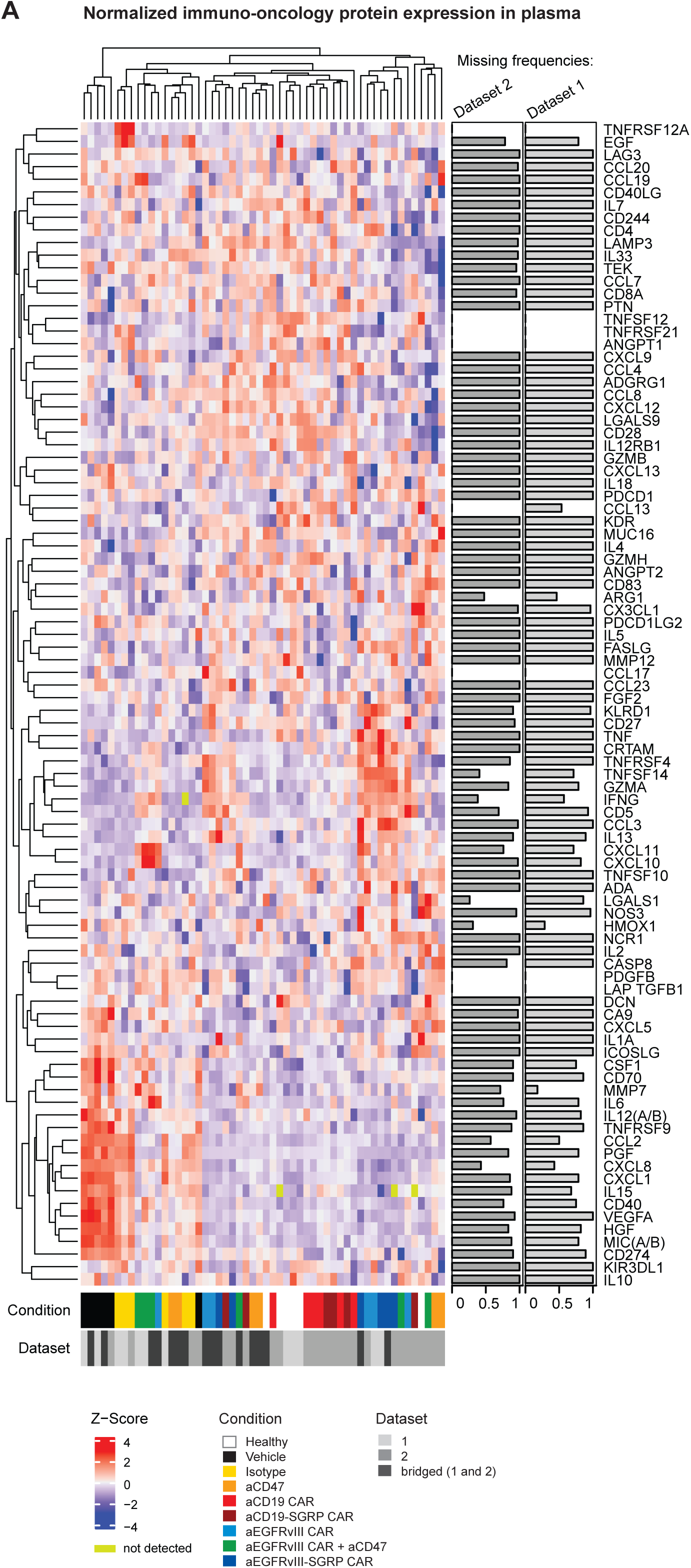
Clustered heatmap of immune-targeted post-therapeutic plasma proteomics. **A,** Clustered heatmap of immuno-oncology-targeted post-therapy plasma proteomics showing the relative expression of proteins between all treatment groups; the analysis included samples from complementary datasets which were bridged and normalized (from lighter to darker grey: dataset 1, dataset 2, bridge samples). Individual protein expression across the datasets are depicted on the y-axis. Each cell represents the Z-score of all the measurements in this row. Data are clustered according to Euclidean distance. The bar plot on the right shows the proportion of measurements under the limit of detection (LOD) for each protein in each dataset.

**Figure S6.**
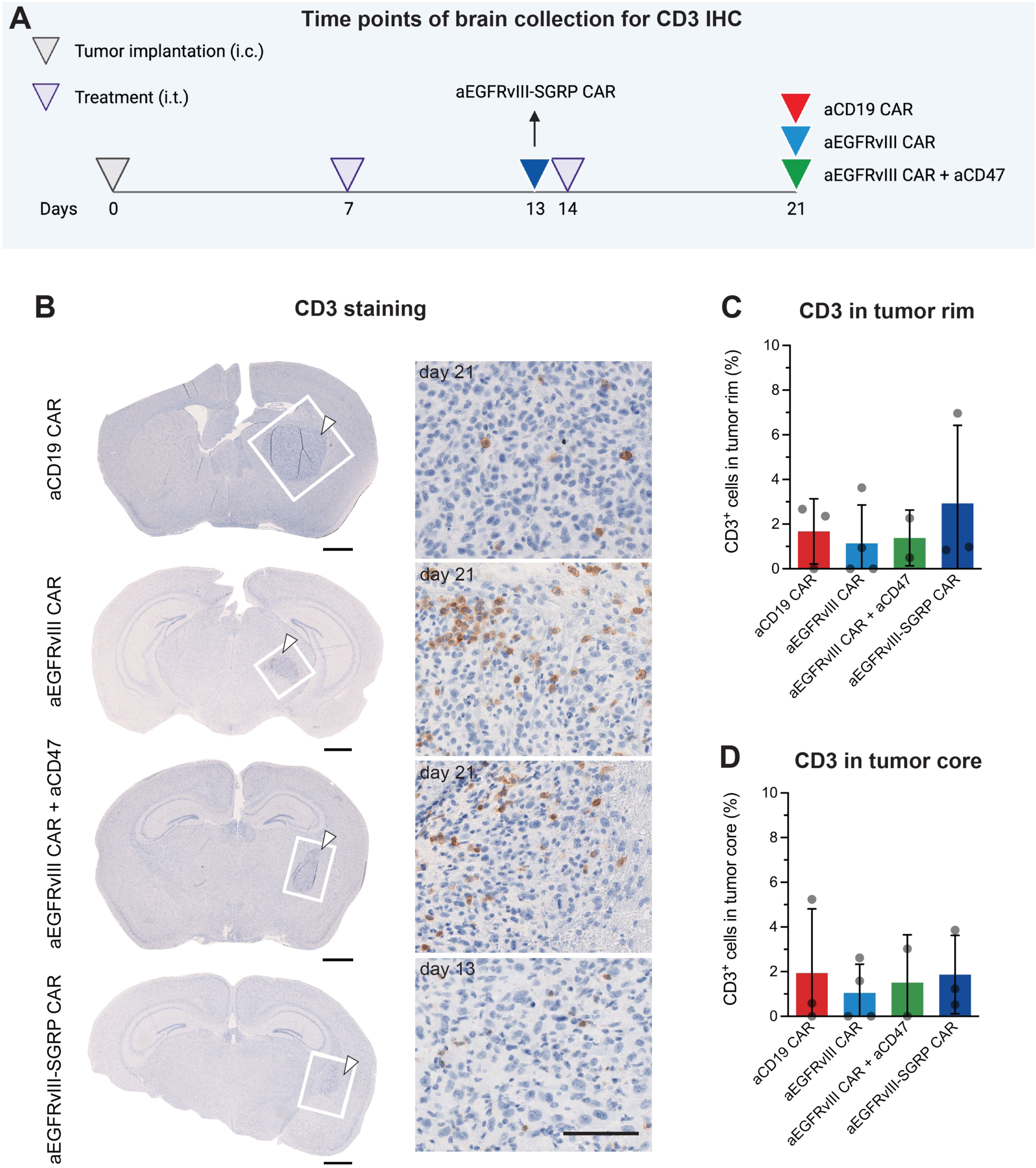
CD3 immunohistochemistry of intermediate time point, post-therapy brains. **A,** Brain collection intermediate post-therapeutic time points for conventional immunohistochemistry per experimental condition. **B,** Representative IHC micrographs of mouse brain sections per therapeutic condition at the indicated time points showing human CD3 staining of grafted T CAR T cells within the tumor core, tumor rim and adjacent brain tissue; *Left column*: overview of tumor-burdened brain sections, scale bar: 1 mm; *Right column*: close-up according to the inserts at the tumor-brain interface, scale bar: 100 μm. **C,** Quantification of CD3^+^ cells in the tumor rim. **D,** Quantification of CD3^+^ cells in the tumor core.

**Figure S7.**
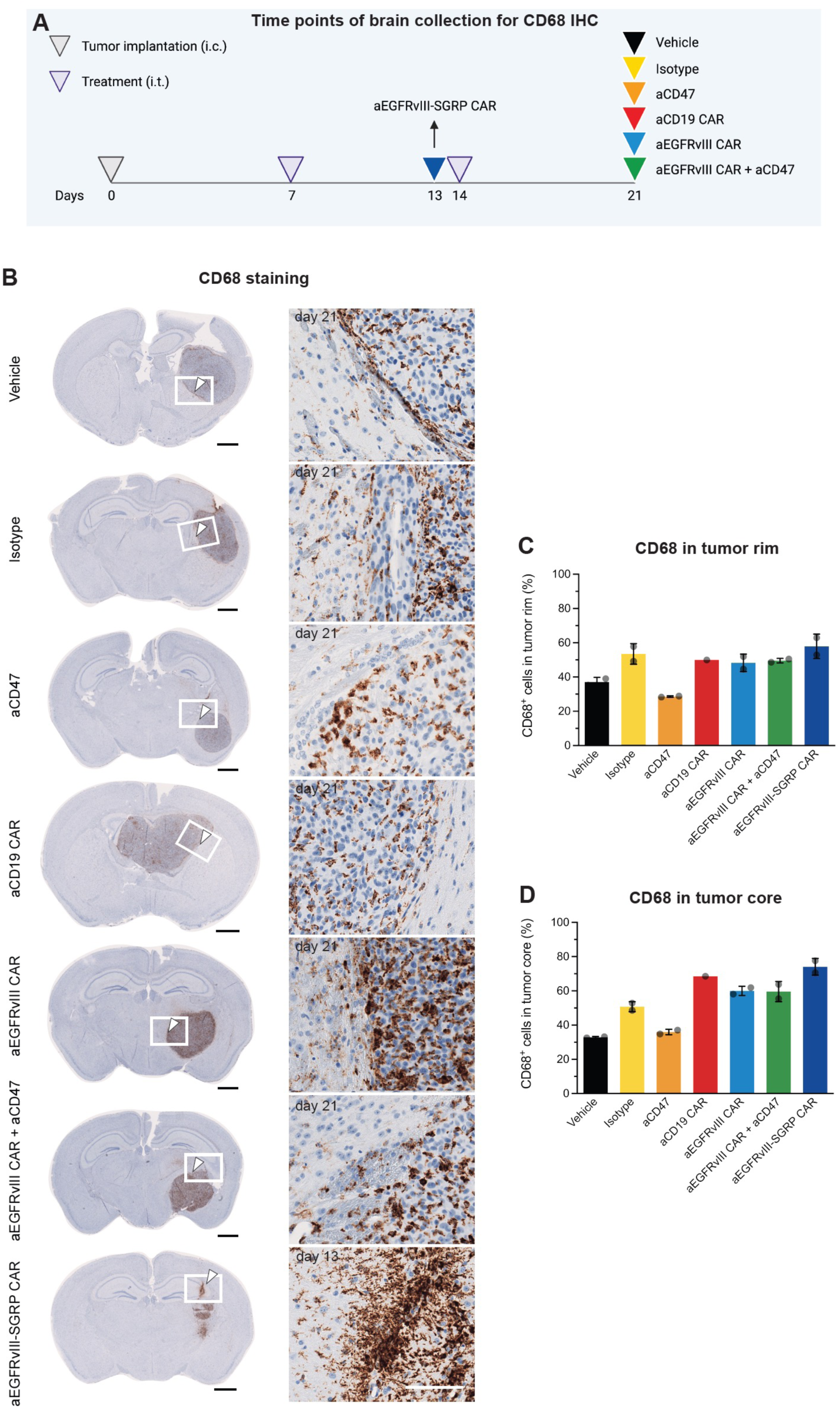
CD68 immunohistochemistry of intermediate time point, post-therapy brains. **A,** Brain collection intermediate post-therapeutic time points for conventional immunohistochemistry per experimental condition. **B,** Representative IHC micrographs of mouse brain sections sections per therapeutic condition at the indicated time points showing mouse CD68 staining of infiltrating myeloid cells within the tumor core, tumor rim and adjacent brain tissue; *Left column*: overview of tumor-burdened brain sections, scale bar: 1 mm; *Right column*: close-up according to the inserts at the tumor-brain interface, scale bar: 100 μm. **C,** Quantification of CD68^+^ cells in the tumor rim. **D,** Quantification of CD68^+^ cells in the tumor core.

**Figure S8.**
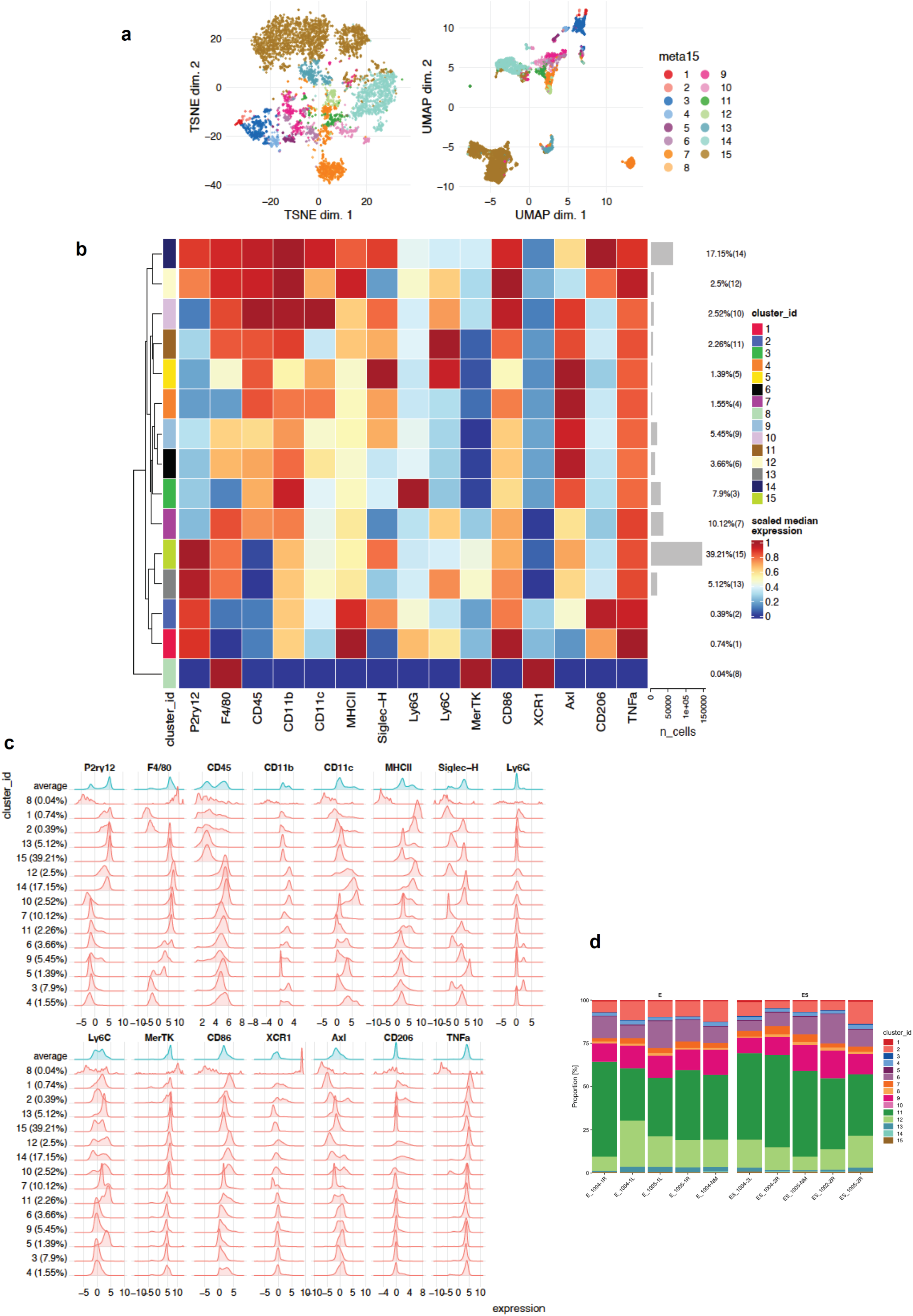
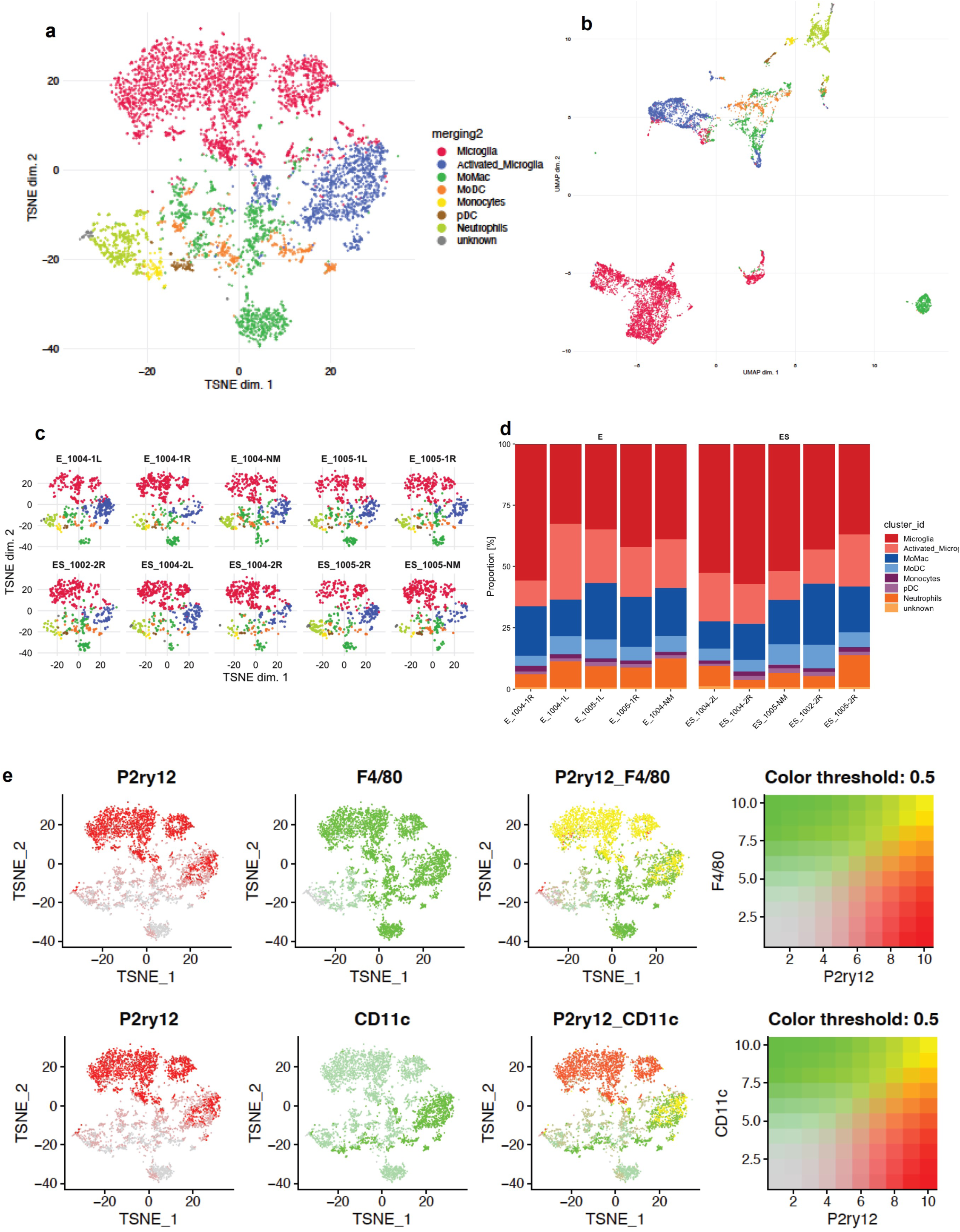
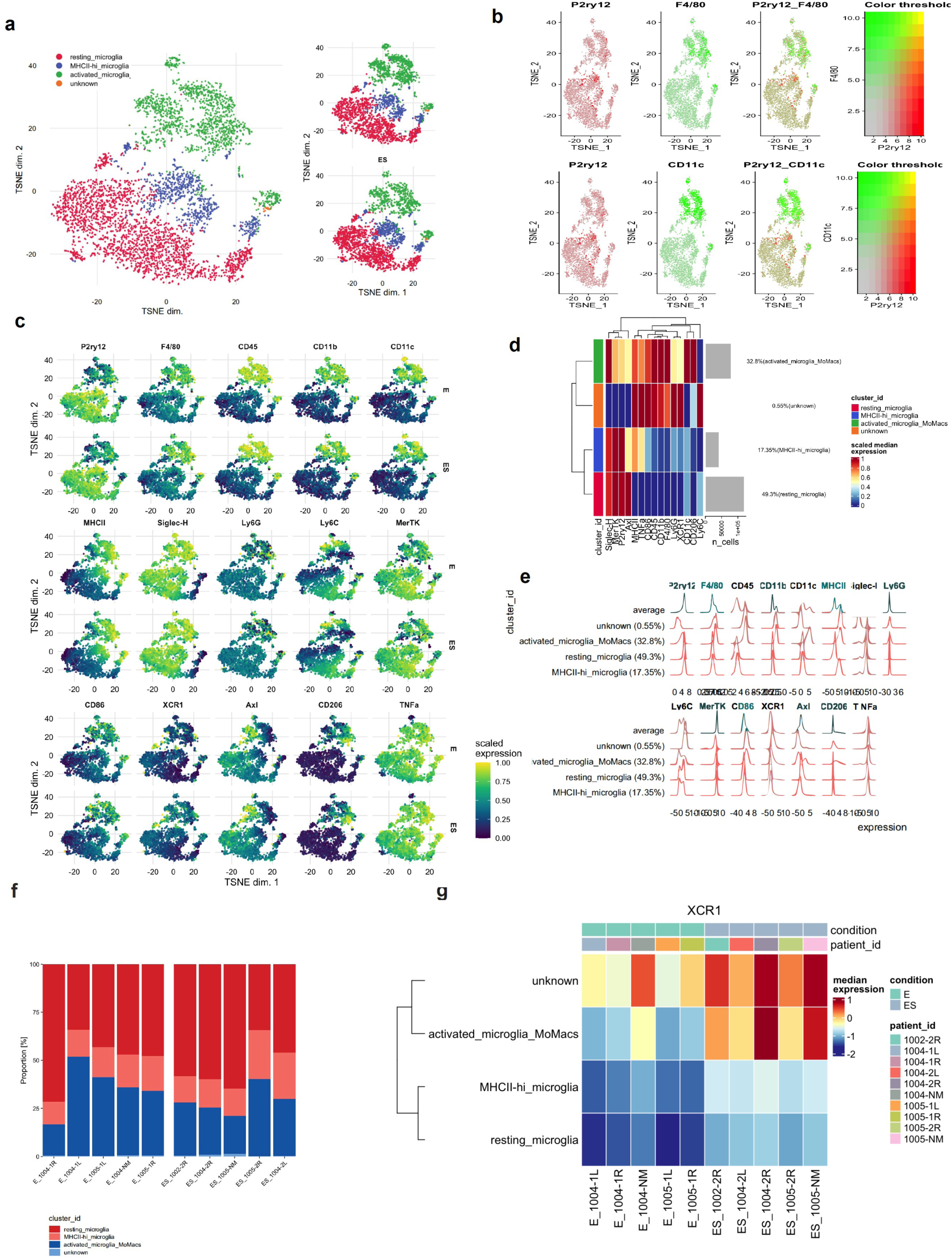
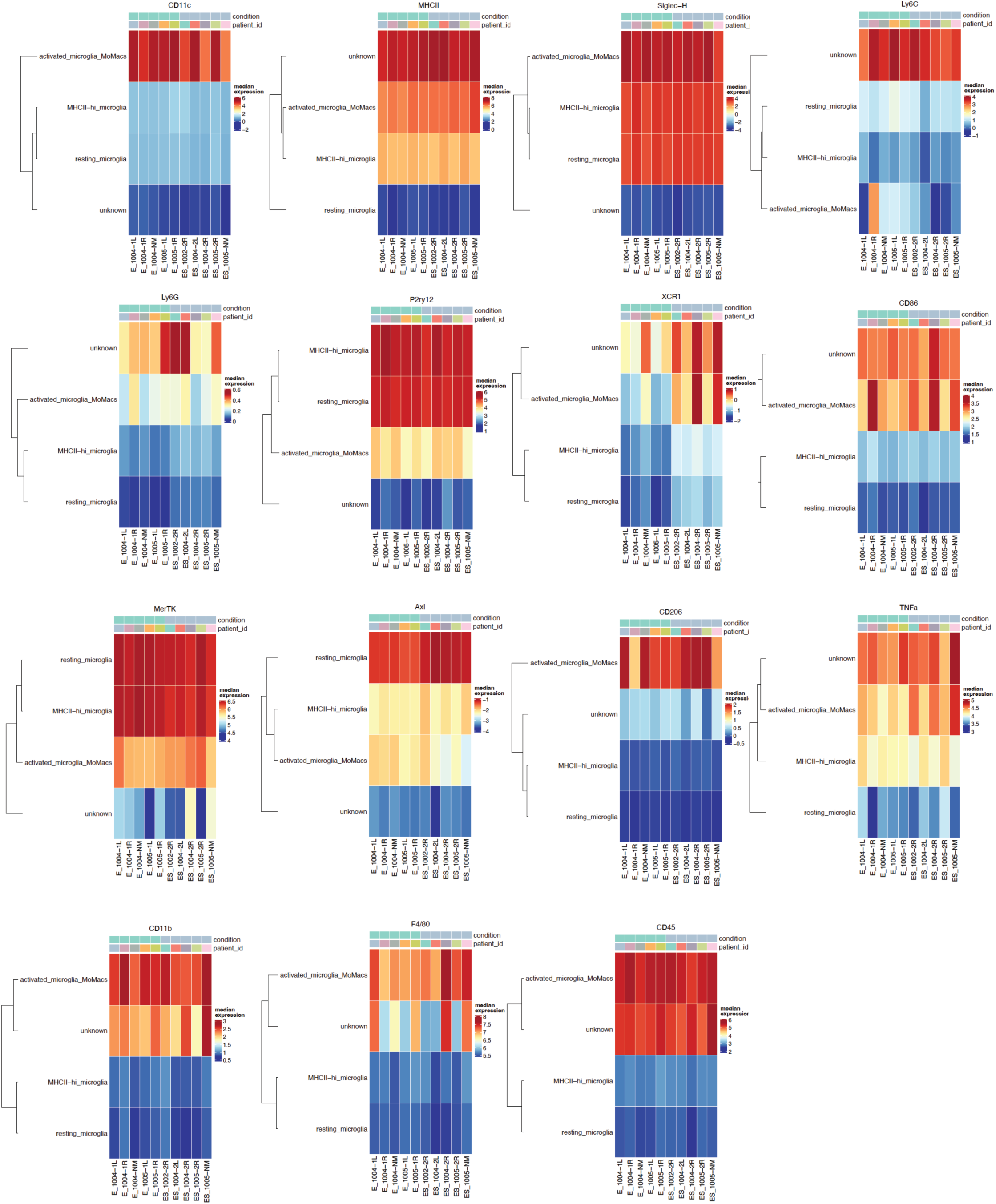
FlowSOM analysis pipeline for spectral flow cytometry. Page 1: **A,** TSNE (left) and UMAP (right) plots depicting the initial 15 clusters within the CD45^+^ pre-gated population. **B,** clustered heatmap of scaled median marker expression per cluster. The number of cells per cluster is indicated on the right y-axis. **C,** Histograms of marker expression per cluster. d) Frequency of clusters in aEGFRvIII CAR-treated animals (E, n = 5 animals), and aEGFRvIII-SGRP CAR-treated animals (ES, n = 5 animals). **Page 2: A,** TSNE (left) and **B,** UMAP (right) plots after merging and annotation of initial clustering. 8 clusters (microglia, activated microglia, MoMac, MoDC, monocytes, pDC, neutrophils, and unknowns) were identified. **C,** Individual TSNEs per experimental condition and animal (E = aEGFRvIII CAR, ES = aEGFRvIII-SGRP CAR treatment). **D,** Stacked bar plots representing cluster frequencies per condition. **E,** Overlay of P2ry12/F4/80 (upper panel) and P2ry12/CD11c on the TSNE plot from **A,** displaying differential contribution of these markers to the microglia subclusters. **Page 3: A,** left panel: TSNE plot of microglia-specific sub clustering, resulting in 3 microglia populations: resting microglia, MHCII-high microglia and activated microglia. Right upper panel: merged TSNE plot for aEGFRvIII CAR-treated animals; right upper panel: TSNE plot for aEGFRvIII-SGRP CAR-treated animals. **B,** Overlay of P2ry12 / F4/80 (upper panel) and P2ry12 / CD11c on the TSNE plot from a) displaying differential contribution of these markers to the microglia subclusters. **C,** individual marker overlay (scaled expression according to color code) on microglia-subclustered t SNE plots. **D,** scaled median expression clustered heatmap of microglia subpopulations. Right y-axis: amount of cells per cluster. **E,** histograms displaying individual marker expression per microglia subcluster. **F,** stacked bar plots displaying the frequency distribution of microglia subpopulations per individual experimental animal. **G,** median expression heatmap displaying XCR1 expression levels in microglia subpopulation per experimental condition/individual animal. **Page 4:** Overview of median expression heatmaps per marker assessed per microglia subpopulations.

**Figure S9.**
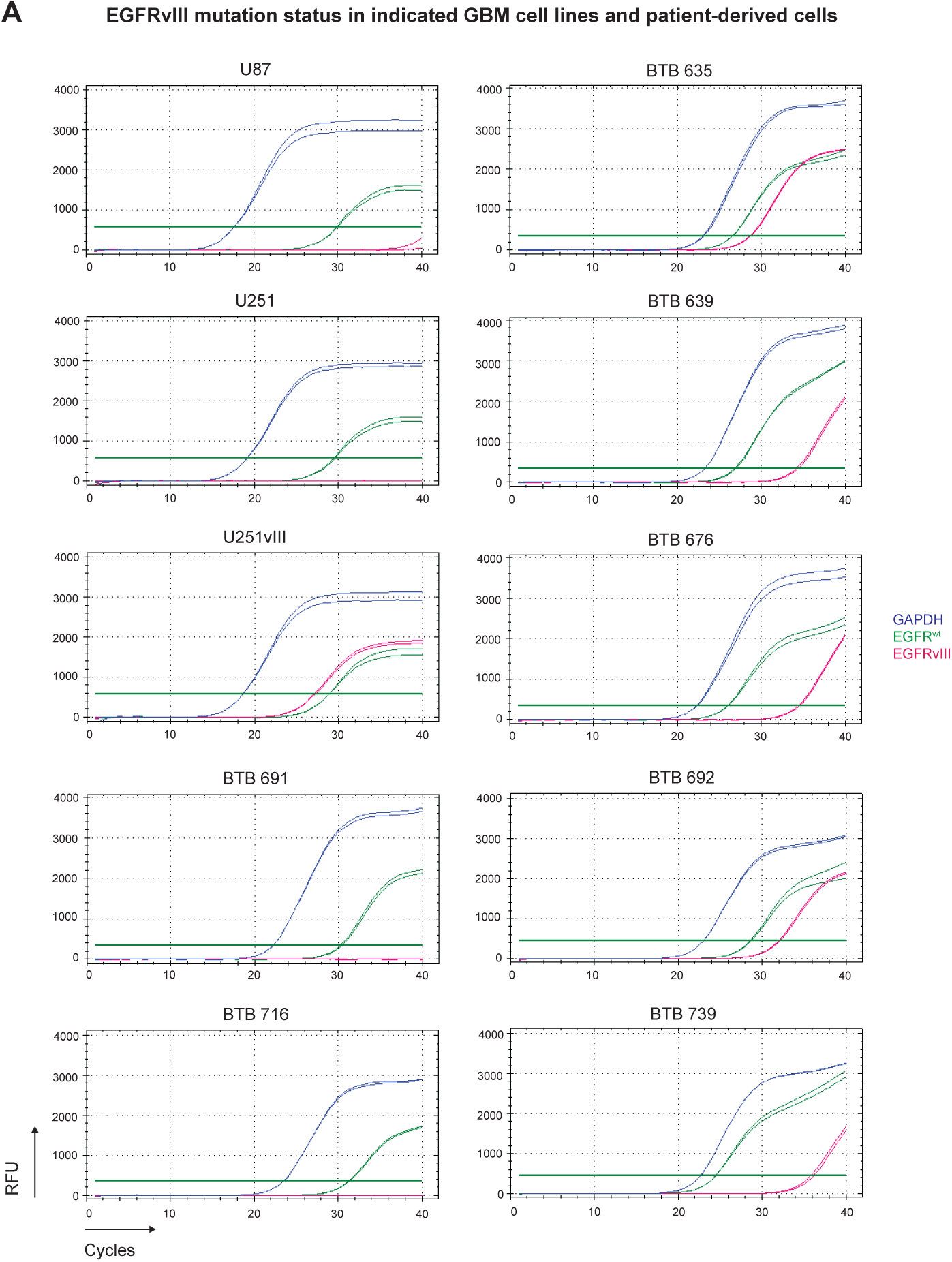
EGFR^wt^ and EGFRvIII screening in GBM cell lines and patient-derived GBM single-cell suspensions. **A,** Detection of EGFR^wt^ and EGFRvIII by real time quantitative PCR (RT-qPCR) in indicated patient-derived single-cell suspensions, showing EGFRvIII positivity in BTB 635, BTB 639, BTB 676, BTB 692 and BTB 739, and no EGFRvIII detection in BTB 691 and BTB 716. GBM cell lines were used as controls for EGFRvIII expression: EGFRvIII^-^ U87 and U251; EGFRvIII^+^ U251vIII. All GBM cell lines and patient-derived samples were positive for EGFR^wt^. RFU: relative fluorescent units.

**Figure S10.**
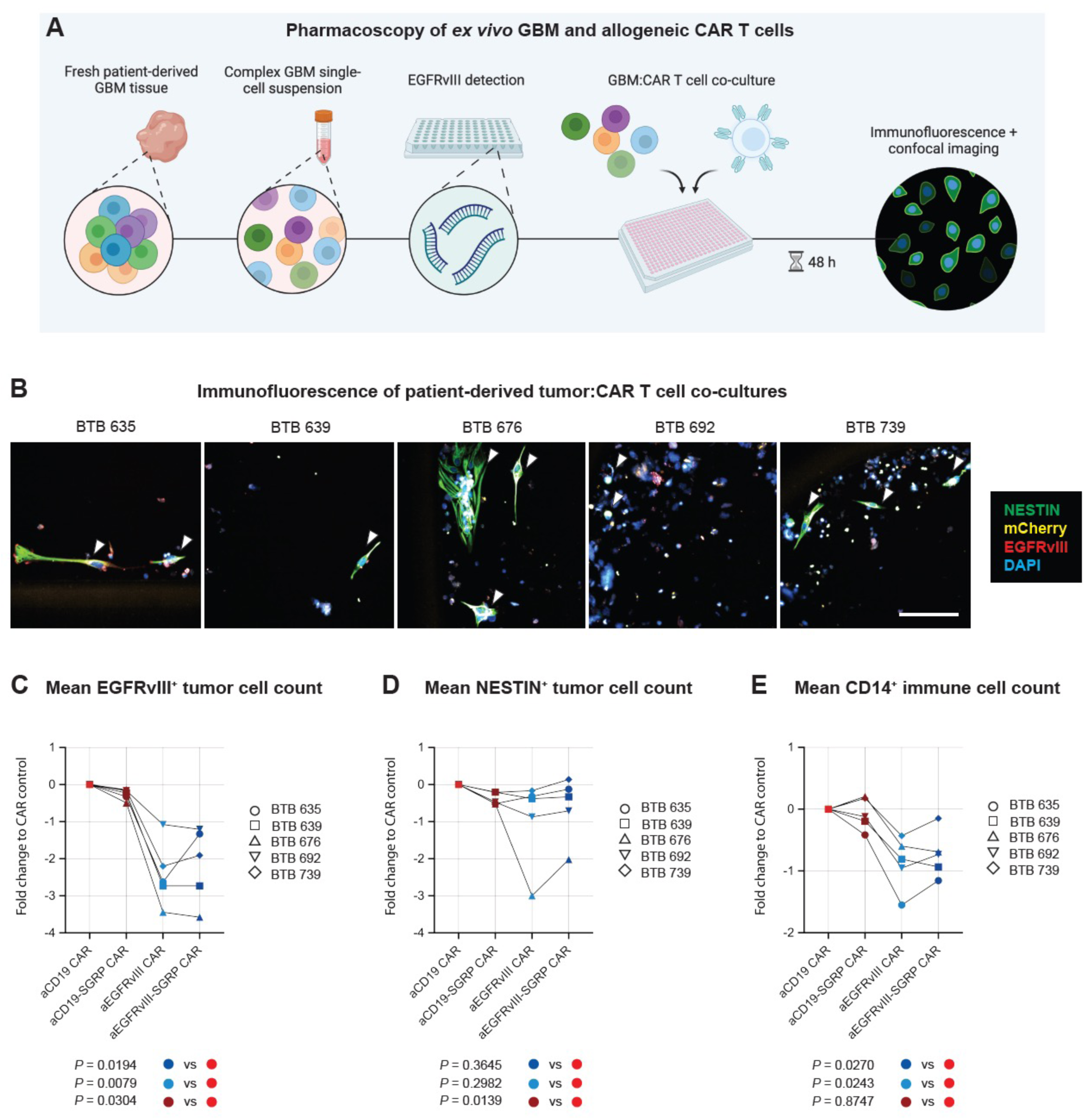
Pharmacoscopy of GBM-CAR co-cultures. **A,** Experimental setup. Batched, frozen single-cell suspensions from GBM patients (tumor center) were thawed and plated at equal numbers into 384 well plates. Beforehand, EGFRvIII status was determined on RNA extracts of matching samples. Single-cell suspensions were co-cultured with aEGFRvIII, aEGFRvIII-SGRP, aCD19 or aCD19-SGRP CAR T cells for 48 h, fixed, stained and imaged via confocal microscopy. **B,** Exemplary immunofluorescence readouts of co-cultures of CAR T cells with single-cell suspensions derived from 5 GBM patients; Scale: 100 µm. **C,** Mean EGFRvIII^+^ tumor cell count displayed as fold change relative to the aCD19 CAR control. **D,** Mean Nestin^+^ tumor cell count displayed as fold change relative to the aCD19 CAR control. **E,** Mean CD14^+^ tumor cell count displayed as fold change relative to the aCD19 CAR control. **C,** to **E,** Statistics: One-way ANOVA with multiple comparisons correction. Source data: **Table S11**.

## Author Contributions

Conceptualization: T.A.M. and G.H.

Data curation: T.A.M., N.T., D.K., S.H., E.M.B., R.W., A.Bu., A.Be. and A.H.

Formal Analysis: T.A.M., D.K., S.H., E.M.B. and A.Bu.

Funding acquisition: G.H.

Investigation: T.A.M., N.T., D.K., R.W., M.-F.R., A.Bu., M.M., A.G., M.M.E., A.H., T.S., H.M., P.S. and I.A.

Methodology: T.A.M., N.T., D.K., S.H., E.M.B., R.W., M.-F.R., A.Bu., A.Be. and J.-L.B.

Project administration: T.A.M. and G.H.

Resources: B.S., T.W., H.L. and G.H.

Software: T.A.M., S.H. and E.M.B.

Supervision: G.H.

Validation: T.A.M., N.T., D.K., S.H., E.M.B., R.W., A.Bu., A.Be., M.M. and A.G.

Visualization: T.A.M., S.H., E.M.B., A.Bu., A.G. and G.H.

Writing – original draft: T.A.M. and G.H.

Writing – review & editing: All authors.

## Acknowledgments

We are grateful to the patients and their families for their consent to donate tissue to our Brain Tumor Biobank. We thank the Neurosurgery Department and the Blood Donation Centre at the University Hospital Basel and all study participants for their invaluable role in this project. We thank the Animal Facility of the Department of Biomedicine at the University of Basel and all animals used in this study. We thank all the staff of the core facilities for Flow Cytometry, Histology, Microscopy (Department of Biomedicine) and Proteomics (Biozentrum) at the University of Basel. We are particularly grateful to Dr. Jelena Markovic-Djuric, Dr. Diego Calabrese, Dr. Michael Abanto, Dr. Alexander Schmidt and Dr. Katarzyna Buczak for their excellent assistance and technical input. The following materials were kindly provided by our collaborators: BS153 cell line was provided by Prof. Alfred Zippelius (University of Basel, Switzerland); HEK293T cell line was provided by Prof. Mohamed Bentires-Alj (University of Basel, Switzerland); NSC197 cell line was provided by Prof. Sheila Singh (McMaster University, Canada); Raji cell line was provided by Prof. Heinz Läubli (University of Basel, Switzerland); U87 cell line was provided by Prof. Luigi Mariani (University Hospital Basel, Switzerland); U251 and U251vIII cell lines were provided by Prof. Sebastian Kobold (Ludwig Maximilian University of Munich, Germany).

## Data Access

All data in this study are available within the manuscript and the associated supplementary materials. The CAR secretome dataset will be deposited in the Proteomics Indentifications Database (PRIDE). Proprietary scripts will be made available via GitHub.

