## Supplementary material for "Enhancing anti-EGFRvIII CAR T cell therapy against glioblastoma with a paracrine SIRPγ-derived CD47 blocker": Table S6

Sex: _______________

Cage: _______________

Injection date: 202	/	/	Mouse strain: _______________	Project/cells injected: _________________________________________

| **Mouse** | **Date** |
| --- | --- |
|  | Intervention |
|  | Weight |
|  | Tumor |
|  | Score |
|  | Intervention |
|  | Weight |
|  | Tumor |
|  | Score |
|  | Intervention |
|  | Weight |
|  | Tumor |
|  | Score |
|  | Intervention |
|  | Weight |
|  | Tumor |
|  | Score |
|  | Intervention |
|  | Weight |
|  | Tumor |
|  | Score |
|  | Intervention |
|  | Weight |
|  | Tumor |
|  | Score |

Administered **drug(s)**	Name, dose, and route

Bioluminescence **imaging**	Tumor detected (+/-)

Euthanasia **score**	0-4: Normal/mild distress, monitor 1-2x/week; 5-9: Moderate distress, monitor daily; 10-15: Severe distress, euthanasia

**Euthanasia score**

| **Type** | **Score** | **Observations** | **Date** |
| --- | --- | --- | --- |
|  |  |  | **Animal** |
| Appearance | 0  1  2  3 | Normal posture and smooth fur  Hunched posture OR slightly ruffled fur  Hunched posture AND slightly ruffled fur  Hunched posture AND completely ruffled fur | |
| Weight | 0  1  2  3 | No loss or loss of less than 10% of body weight  Loss of 10% of body weight  Loss of 15% of body weight  Loss of 20% of body weight | |
| Signs | 0  1  2  3 | Normal respiratory rate and pattern  Slight changes, increased respiratory rate only  Increased rate with abdominal breathing  Decreased rate with abdominal breathing | |
| Movement | 0  1  2  3 | Moves normally around the cage  Moves slowly around the cage  Moves only when touched  Does not move OR self-mutilation | |
| Behavior | 0  1  2  3 | Normal (moves when cage is disturbed, runs from hand)  Attenuated or exaggerated responses (moves away briskly)  Moderate change (moves away slowly)  Reacts violently OR unresponsive | |
| Euthanasia | 3 is scored in any one category OR at least 10 is scored across different categories | | |
