## Supplementary material for "Enhancing anti-EGFRvIII CAR T cell therapy against glioblastoma with a paracrine SIRPγ-derived CD47 blocker": Table S10

| **EGFR WT qPCR primer sequence (5'->3')** | **Template strand** | **Length** | **Start** | **Stop** | **Tm** | **GC%** | **Self complementarity** | **Self 3’ complementarity** |
| --- | --- | --- | --- | --- | --- | --- | --- | --- |
| **Forward primer** | TATGTCCTCATTGCCCTCAACA | 22 | 262 | 283 | 59.42 | 45.45 | 3.00 | 0.00 |
| **Reverse primer** | CTGATGATCTGCAGGTTTTCCA | 22 | 323 | 302 | 58.65 | 45.45 | 6.00 | 4.00 |
| **Product length** | 62 |  |  |  |  |  |  |  |

| **EGFRvIII qPCR primer sequence (5'->3')** | **Template strand** | **Length** | **Start** | **Stop** | **Tm** | **GC%** | **Self complementarity** | **Self 3’ complementarity** |
| --- | --- | --- | --- | --- | --- | --- | --- | --- |
| **Forward primer** | CTGCTGGCTGCGCTCTG | 17 | 40 | 56 | 61.22 | 70.59 | 4.00 | 2.00 |
| **Reverse primer** | GTGATCTGTCACCACATAATTACCTTTC | 28 | 111 | 84 | 60.42 | 39.29 | 6.00 | 0.00 |
| **Product length** | 72 |  |  |  |  |  |  |  |
