## Supplementary protocol 1 for "Enhancing anti-EGFRvIII CAR T cell therapy against glioblastoma with a paracrine SIRPγ-derived CD47 blocker"

### Protocol # 87 : CD3_ab5690r_1-100 (13/12/2019)

**Version: 1** **Validated: No** **Active: Yes**

#### Procedure: RUO DISCOVERY Universal ( v22.00.0000 )

**DISCOVERY ULTRA**

**Ventana Medical Systems, Inc., 1910 Innovation Park Drive Tucson, Arizona USA**

| Step No | Procedure Step |
|---|---|
| 1 | [ version 22 ] |
| 2 | [ For Research Use Only. Not intended for diagnostic purposes. ] |
| 3 | Enable Mixers |
| 4 | Warmup Slide to 37 Deg C |
| 5 | [ Delay refers to a time delayed start: Select time until run start ] |
| 6 | [ RECOMMENDED: Set temperature to 60°C and time to 8 minutes ] |
| 7 | Disable Mixers |
| 8 | Warmup Slide to [60 Deg C], and Incubate for [20 Minutes] ( Baking ) |
| 9 | Disable Slide Heater |
| 10 | Enable Mixers |
| 11 | Disable Mixers |
| 12 | Warmup Slide to 58 Deg C, and Incubate for 4 Minutes |
| 13 | Apply CC Coverslip Long |
| 14 | Incubate for 8 Minutes |
| 15 | Apply CC Coverslip Long |
| 16 | Incubate for 8 Minutes |
| 17 | Apply CC Coverslip Long |
| 18 | Incubate for 8 Minutes |
| 19 | Apply EZPrep Volume Adjust |
| 20 | Enable Mixers |
| 21 | [ RECOMMENDED: Set temperature to 69°C and time to 8 minutes for each cycle ] |
| 22 | [ Depar Cycle 1 ] |
| 23 | Warmup Slide to [69 Deg C], and Incubate for [8 Minutes] ( Cycle 1 ) |
| 24 | Rinse Slide With EZ Prep |
| 25 | Apply Coverslip |
| 26 | Incubate for 4 Minutes |
| 27 | Apply EZPrep Volume Adjust |
| 28 | [ Depar Cycle 2 ] |
| 29 | Incubate for [8 Minutes] ( Cycle 2 ) |
| 30 | Rinse Slide With EZ Prep |
| 31 | Apply Coverslip |
| 32 | Incubate for 4 Minutes |
| 33 | Apply EZPrep Volume Adjust |
| 34 | [ Depar Cycle 3 ] |
| 35 | Incubate for [8 Minutes] ( Cycle 3 ) |
| 36 | Rinse Slide With EZ Prep |
| 37 | Apply Depar Volume Adjust |
| 38 | Apply Coverslip |
| 39 | Warmup Slide to 37 Deg C |
| 40 | Rinse Slide With EZ Prep |
| 41 | Apply Long Cell Conditioner #1 |
| 42 | Apply CC Coverslip Long |
| 43 | Warmup Slide to [95 Deg C], and Incubate for 4 Minutes ( Cell Conditioner #1 ) |
| 44 | Incubate for 4 Minutes |

| Step No | Procedure Step |
|---|---|
| 45 | Incubate for 8 Minutes |
| 46 | Apply Cell Conditioner #1 |
| 47 | Apply CC Medium Coverslip No BB |
| 48 | Incubate for 8 Minutes |
| 49 | Incubate for 8 Minutes |
| 50 | Apply Cell Conditioner #1 |
| 51 | Apply CC Medium Coverslip No BB |
| 52 | Apply Cell Conditioner #1 |
| 53 | Apply CC Medium Coverslip No BB |
| 54 | Apply Cell Conditioner #1 |
| 55 | Apply CC Medium Coverslip No BB |
| 56 | Disable Slide Heater |
| 57 | Apply Cell Conditioner #1 |
| 58 | Apply CC Medium Coverslip No BB |
| 59 | Warmup Slide to 37 Deg C |
| 60 | Rinse Slide With Reaction Buffer |
| 61 | Adjust Slide Volume With Reaction Buffer |
| 62 | Apply Coverslip |
| 63 | [ Select an Inhibitor ] |
| 64 | [ NOTE: Inhibitor CM comes packaged with Chromomap DAB; InhibitorD comes packaged with DABMap ] |
| 65 | [ DISCOVERY Inhibitor is a stand alone product for use with all other HRP substrates ] |
| 66 | [ Inhibitor CM will be applied ] |
| 67 | Rinse Slide With Reaction Buffer |
| 68 | Adjust Slide Volume With Reaction Buffer |
| 69 | Apply Coverslip |
| 70 | Rinse Slide With Reaction Buffer |
| 71 | Adjust Slide Volume With Reaction Buffer |
| 72 | Apply Coverslip |
| 73 | Apply One Drop of Inhibitor CM, and Incubate for [8 Minutes] |
| 74 | Rinse Slide With Reaction Buffer |
| 75 | Adjust Slide Volume With Reaction Buffer |
| 76 | Apply Coverslip |
| 77 | Disable Slide Heater |
| 78 | Disable Mixers |
| 79 | Wait For Button ( Antibody ) |
| 80 | Enable Mixers |
| 81 | Warmup Slide to 37 Deg C |
| 82 | Incubate for 4 Minutes |
| 83 | Rinse Slide With Reaction Buffer |
| 84 | Adjust Slide Volume With Reaction Buffer |
| 85 | Apply Coverslip |
| 86 | Warmup Slide to [37 Deg C] from Very Low Temperatures ( Primary Antibody ) |
| 87 | Hand Apply ( Primary Antibody ), and Incubate for [60 Minutes] |
| 88 | Rinse Slide With Reaction Buffer |

* one drop is one reagent dispense

| Step No | Procedure Step |
|---|---|
| 89 | Adjust Slide Volume With Reaction Buffer |
| 90 | Apply Coverslip |
| 91 | [ Inhibitor Solution will not be applied after the primary ] |
| 92 | Disable Slide Heater |
| 93 | Warmup Slide to 37 Deg C |
| 94 | [ Requires DETECTION dispensers ] |
| 95 | [ These selections may be used for haptenated linking antibodies ] |
| 96 | Rinse Slide With Reaction Buffer |
| 97 | Adjust Slide Volume With Reaction Buffer |
| 98 | Apply Coverslip |
| 99 | Rinse Slide With Reaction Buffer |
| 100 | Adjust Slide Volume With Reaction Buffer |
| 101 | Apply Coverslip |
| 102 | Rinse Slide With Reaction Buffer |
| 103 | Adjust Slide Volume With Reaction Buffer |
| 104 | Apply Coverslip |
| 105 | Incubate for 4 Minutes |
| 106 | Warmup Slide to [37 Deg C] from Very Low Temperatures ( 2nd Antibody ) |
| 107 | Apply One Drop of [DETECTION 1] ( Detection #1 ), and Incubate for [0 Hr 32 Min] |
| 108 | Rinse Slide With Reaction Buffer |
| 109 | Adjust Slide Volume With Reaction Buffer |
| 110 | Apply Coverslip |
| 111 | Disable Slide Heater |
| 112 | Warmup Slide to 37 Deg C |
| 113 | Rinse Slide With Reaction Buffer |
| 114 | Adjust Slide Volume With Reaction Buffer |
| 115 | Apply Coverslip |
| 116 | Apply One Drop of H2O2 CM, and Incubate for 4 Minutes |
| 117 | Apply One Drop of DAB CM, and Incubate for 8 Minutes |
| 118 | Rinse Slide With Reaction Buffer |
| 119 | Adjust Slide Volume With Reaction Buffer |
| 120 | Apply One Drop of Copper CM, Apply Coverslip, and Incubate for 4 Minutes |
| 121 | Rinse Slide With Reaction Buffer |
| 122 | Adjust Slide Volume With Reaction Buffer |
| 123 | Apply Coverslip |
| 124 | Rinse Slide With Reaction Buffer |
| 125 | Adjust Slide Volume With Reaction Buffer |
| 126 | Apply Coverslip |
| 127 | Apply One Drop of [HEMATOXYLIN II] ( Counterstain ), and Incubate for [8 Minutes] |
| 128 | Rinse Slide With Reaction Buffer |
| 129 | Adjust Slide Volume With Reaction Buffer |
| 130 | Apply Coverslip |
| 131 | Rinse Slide With Reaction Buffer |
| 132 | Adjust Slide Volume With Reaction Buffer |

* one drop is one reagent dispense

| Step No | Procedure Step |
|---|---|
| 133 | Apply Coverslip |
| 134 | Apply One Drop of [BLUING REAGENT] ( Post Counterstain ), and Incubate for [8 Minutes] |
| 135 | Rinse Slide With Reaction Buffer |
| 136 | Adjust Slide Volume With Reaction Buffer |
| 137 | Apply Coverslip |
